# Lysosomal microautophagic pathway links proteostasis to protein aggregation disease

**DOI:** 10.1101/2020.08.11.245688

**Authors:** Yuuki Fujiwara, Chihana Kabuta, Viorica Raluca Contu, Megumu Ogawa, Ryohei Sakai, Hiromi Fujita, Hisae Kikuchi, Mari Suzuki, Ikuko Koyama-Honda, Michio Inoue, Yasushi Oya, Yukiko U. Inoue, Takayoshi Inoue, Ryosuke Takahashi, Gen Matsumoto, Kazuaki Matoba, Nobuo N. Noda, Ichizo Nishino, Keiji Wada, Toyoshi Fujimoto, Satoru Noguchi, Tomohiro Kabuta

## Abstract

Constitutive lysosomal degradation of cytosolic proteins is fundamental to proteostasis, but how these substrates are delivered into the lysosomal lumen remains incompletely understood. In mammals, the contribution of microautophagy to basal lysosomal protein degradation and the molecular machinery that mediates this process remain poorly defined. Here we identify a mammalian microautophagic pathway that is mediated by the lysosomal membrane protein SIDT2 and directly delivers cytosolic proteins, including disease-associated aggregation-prone proteins, into the lysosomal lumen for degradation. This pathway functions independently of macroautophagy and chaperone-mediated autophagy. Reconstitution and imaging analyses showed that SIDT2 mediates direct lysosomal uptake and subsequent degradation of cytosolic proteins and increases the prevalence and size of intraluminal vesicles, with the resulting vesicles exhibiting a topology consistent with inward sequestration of cytosol. A conserved catalytic serine residue was required for SIDT2 ATPase activity *in vitro* and was also essential for these increases in prevalence and size of intraluminal vesicles and for protein degradation. Loss of SIDT2 impaired lysosomal substrate uptake without altering luminal acidification or proteolytic activity. We further identified a dominant-negative SIDT2 mutation in familial neuromyopathy with rimmed vacuoles. Both *Sidt2*-deficient mice and knock-in mice carrying the patient mutation recapitulated key pathological features, including accumulation of aggregation-prone proteins and rimmed vacuole-like lesions. These findings establish SIDT2-dependent microautophagy as a pathway for basal lysosomal protein degradation and show that its failure can underlie human protein aggregation disease.

---

Biological homeostasis depends on the cyclic synthesis and degradation of cellular components. The regulated degradation of biomacromolecules is essential for maintaining the intracellular environment. Dysfunction in these degradation pathways is associated with various diseases, such as neurodegenerative and neuromuscular disorders. Lysosomes are the principal organelles involved in the degradation of macromolecules. The cytosolic components are delivered into lysosomes and degraded during autophagy^1,2^. Various cellular components are processed via distinct autophagic pathways^1,2^, including macroautophagy, chaperone-mediated autophagy (CMA), and microautophagy. The best-characterized autophagic pathway is macroautophagy, where various cellular components are sequestered by autophagosomes, double-membraned vesicles, and delivered to lysosomes. In CMA, chaperones assist the direct import of specific groups of proteins into lysosomes, whereas in microautophagy, cytosolic components are sequestered into lysosomes or endosomes by invagination. Compared to macroautophagy and CMA, studies on the molecular mechanisms underlying microautophagy and its physiological significance, particularly in mammalian cells, are limited^3^. Although macroautophagy is the most extensively studied autophagic pathway to date, its contribution to overall lysosomal protein degradation is also unclear. For instance, substantial lysosomal protein degradation persists in *Atg5*-deficient cells^4^. In addition, macroautophagy is primarily known as an inducible pathway that responds to stresses such as starvation^1^. In this study, we investigated the autophagic pathway responsible for basal lysosomal protein degradation independently of macroautophagy.

## Results

### Basal lysosomal proteolysis driven by SIDT2

Under nutrient-rich conditions, the lysosomal acid protease inhibitors pepstatin A and E64d (Pep+E64) significantly reduced total degradation of endogenous proteins in both wild-type (WT) mouse embryonic fibroblasts (MEFs) and MEFs deficient in *Atg5* or *Atg13*, two core factors required for macroautophagy (Fig. 1a and Extended Data Fig. 1a). This suggests that lysosomal degradation pathways other than macroautophagy may play an important role in basal protein turnover.

**Fig. 1.**
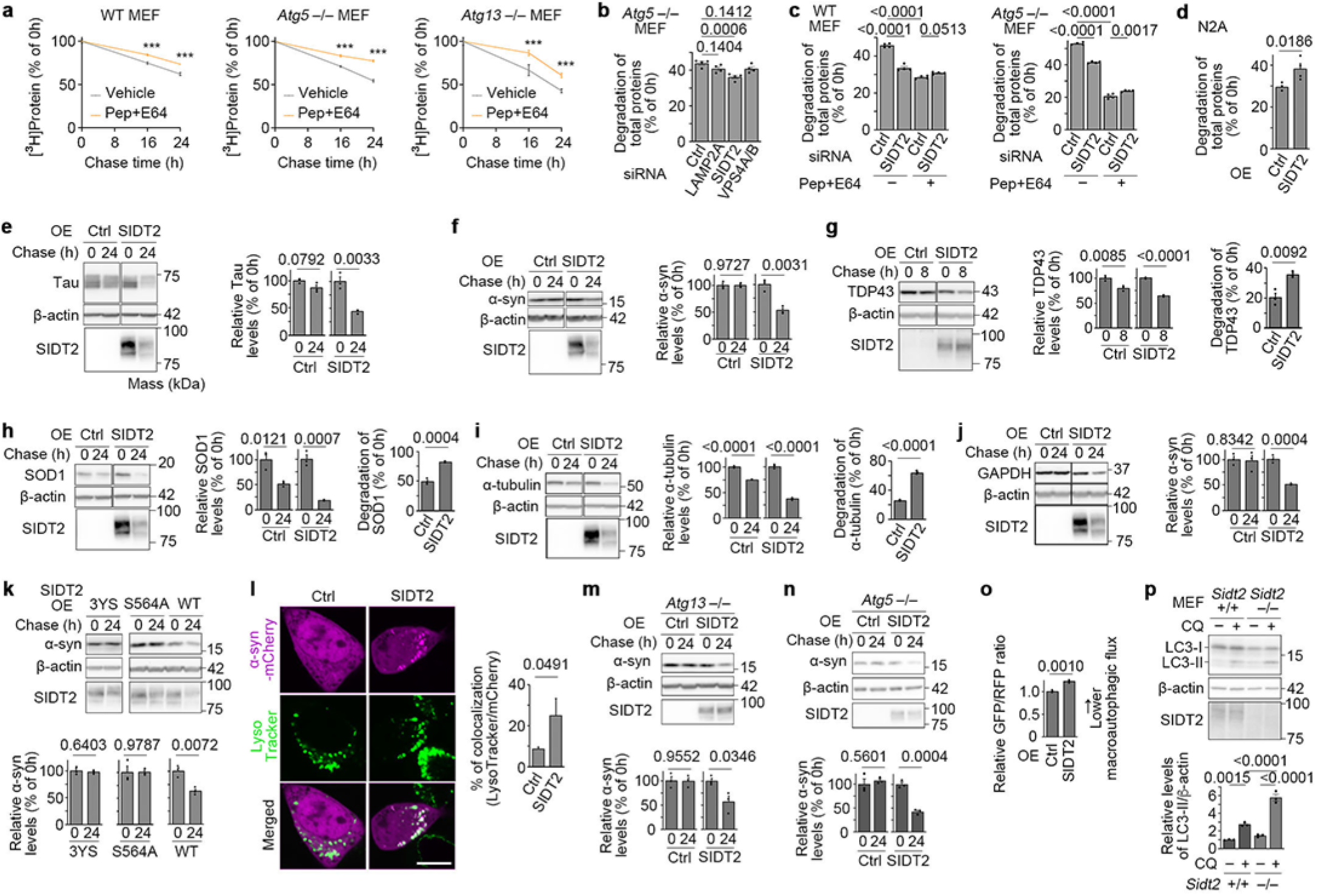
SIDT2 mediates lysosomal uptake and degradation of proteins in cells. (a) Degradation of intracellular proteins in wild-type (WT) (*Atg5*^+/+^), *Atg5*^−/–^, and *Atg13*^−/–^ mouse embryonic fibroblasts (MEFs) under nutrient-rich conditions. Cells were treated with either vehicle (DMSO) or lysosomal protease inhibitors Pepstatin A and E64d (50 µg/mL each) (n = 4). \*\*\**P* ≤ 0.0001 (vs vehicle at each time point). Data are the mean ± s.d. (b) Effect of LAMP2A, SIDT2, or VPS4A knockdown on degradation of radiolabelled total proteins in *Atg5^−/–^*MEFs (n = 4). (c) Effect of SIDT2 knockdown and lysosomal protease inhibitors on the degradation of radiolabelled total protein in WT or *Atg5^−/–^*MEFs (n = 4). (d) Effect of SIDT2 overexpression on the degradation of radiolabelled total protein in N2a cells (n = 4). OE: overexpression. (e–j) Degradation of various cytosolic proteins in N2a cells overexpressing SIDT2, using the Tet-Off system (n = 3). (k) Degradation of α-synuclein protein in N2a cells overexpressing 3YS– or S564A–SIDT2 (n = 3). (l) Localization of α-synuclein–mCherry protein in SIDT2-overexpressing N2a cells (control, n = 14; SIDT2, n = 12). Scale bar, 10 µm. (m, n) Degradation of α-synuclein protein in *Atg13*-knockout (KO) (m) and *Atg5*-KO (n) N2a cells (n = 3). (o) Macroautophagic flux in N2a cells overexpressing SIDT2 (n = 3). The higher GFP/RFP ratio indicates lower activity of macroautophagy. (p) Macroautophagic flux in *Sidt2*^+/+^ and *Sidt2*^−/–^ MEFs, assessed by LC3 immunoblotting with or without chloroquine (CQ) treatment (n = 3).

To investigate this further, we examined the contribution of several autophagic pathways other than macroautophagy in basal protein degradation. While knockdown of LAMP2A and VPS4, required for CMA and a subset of microautophagy, respectively, did not significantly reduce the steady-state protein degradation in *Atg5*-deficient MEFs, knockdown of SIDT2, a lysosomal membrane protein, significantly reduced basal protein degradation levels in both *Atg5*-deficient and WT MEFs (Fig. 1b, c and Extended Data Fig. 1b). This was unexpected because SIDT2 was previously shown to mediate RNautophagy^5^, a macroautophagy-independent process in which lysosomes directly take up and degrade RNA^6^. Furthermore, the suppressive effect of SIDT2 knockdown on protein degradation was abolished by Pep+E64 in both *Atg5*-deficient and WT cells, confirming the involvement of SIDT2 in lysosome-dependent protein degradation (Fig. 1c, Extended Data Fig. 1b, c). Although some reports have suggested that SIDT2 may contribute to macroautophagy activity^7,8^, reduced lysosomal protein degradation upon SIDT2 knockdown even in *Atg5*-deficient cells indicates that SIDT2 functions independently of macroautophagy. Particularly in WT MEFs, SIDT2 knockdown alone reduced protein degradation to a level similar to that observed with Pep+E64 treatment (Fig. 1c), suggesting that the SIDT2-dependent pathway can account for a large proportion of lysosomal degradation of intracellular proteins in these cells. In *Atg5* knockout (KO) MEFs, elevated SIDT2- and macroautophagy-independent lysosomal protein degradation may reflect a compensatory activation of other autophagic pathway(s).

We next sought to analyse the effect of SIDT2 overexpression on protein degradation levels within cells. Considering the relatively high expression levels of endogenous SIDT2 in MEFs, we used Neuro2a (N2a) cells which show substantially lower levels of SIDT2 (Extended Data Fig. 1d). We have previously reported that in N2a cells, overexpression of SIDT2 increases RNautophagy activity^9^. Consistently, overexpression of SIDT2 markedly increased the total degradation of intracellular proteins (Fig. 1d). Furthermore, Tet-Off analyses in N2a, HeLa, and C2C12 cells revealed that overexpression of SIDT2 markedly increased intracellular degradation of both general cytosolic proteins, including superoxide dismutase 1 (SOD1), α-tubulin, and glyceraldehyde 3-phosphate dehydrogenase (GAPDH), and disease-associated aggregation-prone proteins such as tau, α-synuclein and TDP-43 (Fig. 1e–j and Extended Data Fig. 2). Although SIDT2 affected the degradation levels of most cytosolic proteins examined, we selected α-synuclein as a reporter substrate for further analyses because its SIDT2-dependent degradation was readily detectable. The 3YS mutant of SIDT2, which lacks lysosomal targeting signals and is largely excluded from lysosomes^9^, had no effect on the degradation of α-synuclein (Fig. 1k), indicating that localization of SIDT2 to the lysosomal membrane is required for SIDT2-mediated protein degradation. We have previously reported that Ser-564 of SIDT2 is required for the uptake of nucleic acids into lysosomes^5,10^. This substitution completely abolished the SIDT2-dependent increase in intracellular protein degradation (Fig. 1k), suggesting that nucleic acids and proteins may be taken up into lysosomes in a similar SIDT2-dependent manner. SIDT2-dependent accumulation of α-synuclein within lysosomes was observed by fluorescence imaging using fluorescent protein-tagged α-synuclein^11^ (Fig. 1l and Extended Data Fig. 3a, b), providing further evidence of SIDT2-mediated lysosomal protein uptake. We then reanalysed whether the SIDT2-dependent pathway is independent of macroautophagy. First, the SIDT2-dependent increase in protein degradation was observed in cells deficient for either *Atg13* or *Atg5* (Fig. 1m, n and Extended Data Fig. 4a–c). Although some studies have suggested that SIDT2 may be positively associated with the regulation of macroautophagy^7,8^, we found that macroautophagic activity did not increase but rather decreased upon SIDT2 overexpression (Fig. 1o). Consistent with this result, deletion of *Sidt2* in MEFs increased macroautophagic activity (Fig. 1p). Taken together, these results indicate that the SIDT2-dependent pathway in cells is independent of macroautophagy.

### SIDT2 mediates direct lysosomal protein uptake

Since lysosomes directly internalize substrates in autophagic pathways other than macroautophagy, we investigated whether SIDT2 mediates direct lysosomal protein uptake using an in vitro reconstitution assay involving lysosomal preparations^5,6,10,12^ with α-synuclein, tau, and SOD1 proteins as substrates. Lysosomes were co-incubated with substrate proteins (± ATP), and the samples were analysed by western blotting and immunoelectron microscopy (Fig. 2a). All three proteins examined were directly degraded by intact lysosomes only in the presence of ATP (Fig. 2b, c and Extended Data Fig. 5a–c). Because lysosomal luminal hydrolases do not require ATP for their catalytic activity^13^, ATP is likely required for protein uptake, consistent with our previous findings that nucleic acid uptake by lysosomes depends on ATP consumption^14^. Further, the protein degradation was abolished by pre-incubation of lysosome preparations with Pep+E64 (Fig. 2d). In addition, protein degradation was not observed in the extralysosomal solution collected after pre-incubation of lysosome preparations with ATP (Extended Data Fig. 5d). These data indicate that protein degradation occurs in the lysosomal lumen. Lastly, using immunoelectron microscopy, we observed that α-synuclein was directly taken up into the lysosomal lumen (Fig. 2e and Extended Data Fig. 5e–g). Collectively, these data strongly suggest that proteins are directly taken up and degraded in lysosomes.

**Fig. 2.**
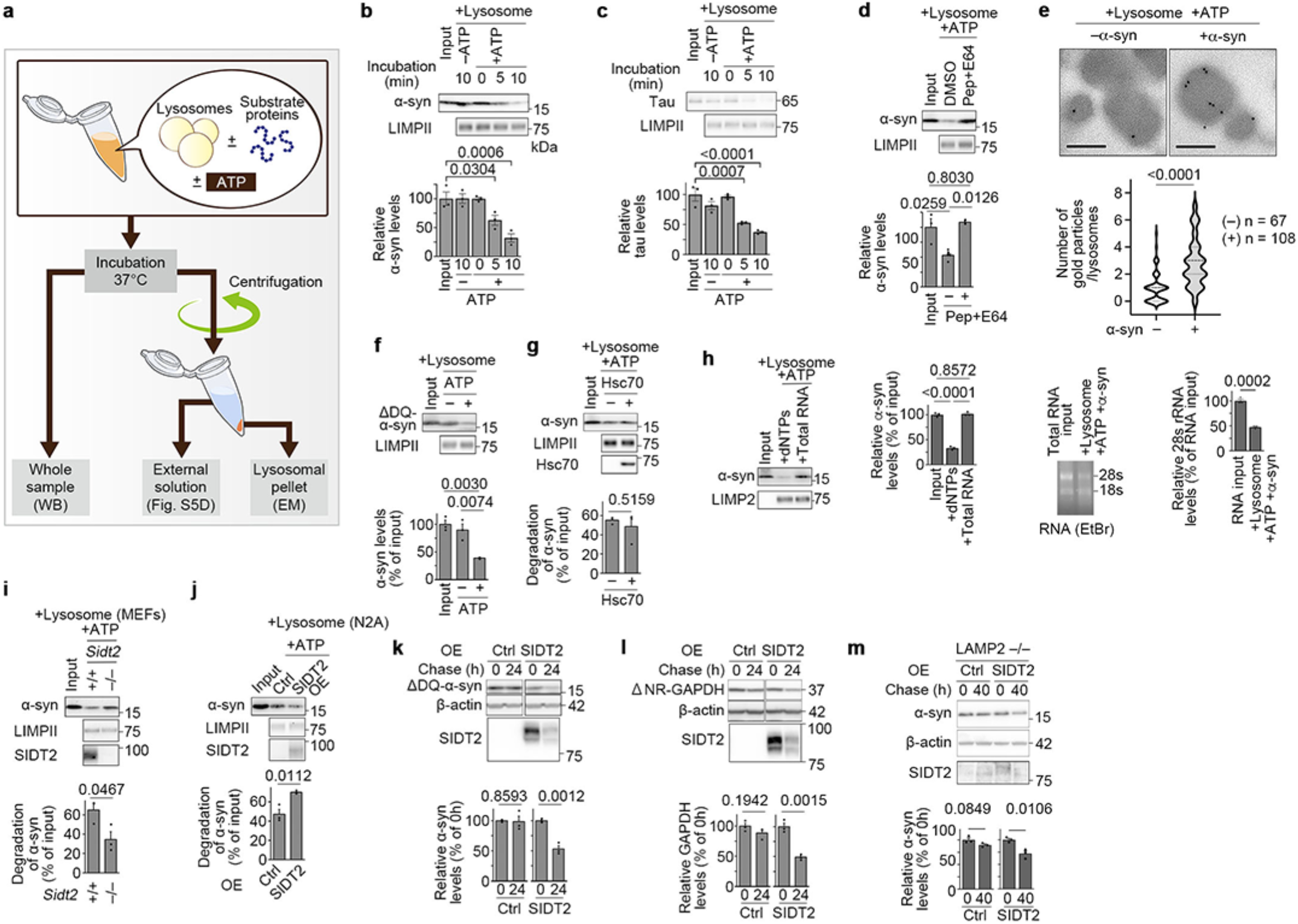
Lysosomal proteolysis through direct uptake of protein into lysosomes via SIDT2. (a) Schematic of an in vitro reconstruction assay using lysosomal preparations. (b, c) Degradation of α-synuclein (b) and tau (c) protein by lysosomes derived from WT MEFs. (d) Degradation of α-synuclein protein by lysosomes derived from WT MEFs preincubated with or without Pep+E64. (e) Immunoelectron microscopy image of α-synuclein protein in lysosomal preparations co-incubated with or without recombinant α-synuclein protein in the presence of ATP. Quantification of gold particles per lysosome is shown below. Scale bars, 200 nm. (f) Degradation of recombinant mutant α-synuclein protein that lacks the KFERQ-like motif by lysosomes. (g) Degradation of recombinant WT α-synuclein protein by lysosomes with or without Hsc70 chaperone. (h) Degradation of α-synuclein by lysosomes in the presence of dNTPs or total RNA (left). Total RNA was degraded under the same conditions (right). (i, j) Degradation of α-synuclein protein by lysosomes derived from *Sidt2*^+/+^ and *Sidt2*^−/–^ MEFs (i) or from SIDT2-overexpressing N2a cells (j). OE: overexpression. (k, l) Degradation of mutant α-synuclein (k) and GAPDH (l) proteins lacking the KFERQ-like motif in N2a cells overexpressing SIDT2, using the Tet-Off system. (m) Degradation of α-synuclein protein in *LAMP2*-KO HeLa cells. For all quantifications except in (e), n = 3.

Among mammalian autophagic pathways, CMA is the only pathway that has been reconstituted using isolated lysosomes and directly mediates protein uptake for degradation. However, in vitro reconstitution of CMA is reported to be restricted to a specific population of CMA-active lysosomes that require enrichment using metrizamide^15^, which was not used in our procedures. We therefore investigated whether the observed lysosomal uptake of proteins is related to CMA. CMA is a selective pathway in which proteins harbouring a KFERQ-like motif are recognized by the chaperone Hsc70^16^ and directly imported into lysosomes via the CMA-specific receptor LAMP2A. α-synuclein contains a KFERQ-like motif required for degradation via CMA^17^. However, ΔDQ α-synuclein, which lacks a KFERQ-like motif and has been reported not to serve as a CMA substrate^17^, was nevertheless degraded by lysosomes in an ATP-dependent manner in vitro (Fig. 2f). Furthermore, unlike in CMA, the addition of Hsc70 did not enhance the degradation of α-synuclein (Fig. 2g). Lastly, we observed that α-synuclein was degraded by lysosomes from LAMP2-deficient HeLa cells, which completely lack the CMA receptor LAMP2A (Extended Data Fig. 5h). These results indicate that the direct uptake and degradation of proteins observed in our in vitro experiments are independent of CMA.

We then analysed whether the direct uptake of protein by lysosomes in vitro is dependent on SIDT2, consistent with its effects in living cells. Because SIDT2 mediates direct lysosomal uptake of nucleic acids^5,10^, we speculated that, if proteins are incorporated into lysosomes via a similar mechanism, addition of RNA may competitively inhibit the protein degradation process by lysosomal preparations. Strikingly, RNA, but not an equal amount of dNTPs, markedly suppressed the degradation of protein by lysosomes (Fig. 2h), suggesting that protein and RNA may share an uptake mechanism into lysosomes. We compared protein degradation levels in lysosomes from *Sidt2*^+/+^ and *Sidt2*^−/–^ MEFs in the presence of ATP and found that degradation was attenuated in *Sidt2*^−/–^ lysosomes (Fig. 2i). *Sidt2*^−/–^ lysosomes showed no apparent morphological changes or alterations in pH and luminal proteolytic activity (Extended Data Fig. 6a–d), suggesting that the attenuated protein degradation is not due to impaired intraluminal degradative capacity or structural changes. Consistent with the result obtained in living cells, lysosomes derived from SIDT2-overexpressing N2a cells showed higher protein degradation activity than control N2a lysosomes (Fig. 2j). In addition, we confirmed that SIDT2 itself does not possess proteolytic activity (Extended Data Fig. 6e). These results indicate that SIDT2 mediates the direct uptake of proteins into lysosomes.

We next examined whether SIDT2-dependent protein degradation is also CMA-independent at the cellular level. ΔDQ α-synuclein^17^ and ΔNR GAPDH proteins, which lack KFERQ-like motifs required for substrate recognition in CMA, were nonetheless degraded in a SIDT2-dependent manner (Fig. 2k, l). Additionally, SIDT2-dependent enhancement of protein degradation persisted in LAMP2-KO HeLa cells (Fig. 2m). Lastly, using the well-established CMA probe assay system^18^, we found that CMA activity is not induced upon SIDT2 overexpression (Extended Data Fig. 7a– c). These results indicate that the SIDT2-dependent uptake and degradation of protein by lysosomes is independent of CMA, both in vitro and at the cellular level.

### Microautophagic protein uptake via SIDT2

Given that SIDT2 mediates direct uptake and degradation of cytosolic proteins by lysosomes independently of macroautophagy and CMA, we next sought to define how substrate proteins enter the lysosomal lumen. In principle, this process could be mediated by a protein-conducting pore/channel or transporter, or by invagination of the lysosomal limiting membrane. Because mCherry fluorescence persisted within lysosomes (Fig. 1l and Extended Data Fig. 3a, b), indicating that substrate proteins at least in part remain folded, we hypothesized that SIDT2-dependent uptake involves microautophagy, rather than channel/transporter-mediated translocation. This hypothesis is consistent with recent cryo-electron microscopy studies showing that the dimeric structure of SIDT2 lacks a discernible nucleic acid conduction pathway large enough to accommodate macromolecules such as RNA but instead contains a membrane-embedded catalytic centre and that SIDT2 exhibits lipid-hydrolytic activity^19,20^. These observations suggested that SIDT2 may promote substrate uptake through lysosomal membrane remodelling, such as microautophagic invagination.

Using super-resolution confocal microscopy, we observed that, in cells overexpressing SIDT2, α-synuclein protein internalized into lysosomes shows punctate intraluminal structures smaller than the lysosomal lumen (Fig. 3a), suggesting that a microautophagic process may occur in the lysosomes (Fig. 3b). However, because these α-synuclein-containing puncta were observed in normal-sized mammalian lysosomes, which are much smaller than the vacuoles in yeast or plants, determining whether they are associated with membrane invaginations or intraluminal vesicle formation remained technically difficult. To visualize such membrane events, we used enlarged endolysosomal compartments induced by the constitutively active Rab5(Q79L) mutant. This approach has been previously used to visualize microautophagy^21^. SIDT2 was localized to the membrane of enlarged Rab5(Q79L)-positive vesicles in N2a cells (Extended Data Fig. 8a). SIDT2 and Rab5(Q79L)-double-positive vesicles are also positive for Lysotracker, which labels acidic compartments, and the Lysotracker signals were eliminated by bafilomycin, a specific inhibitor of v-ATPase (Extended Data Fig. 8a). We also confirmed that the Rab5(Q79L)-positive vesicles are LAMP1-positive (Extended Data Fig. 8b). These data indicate that the enlarged vesicles, which are positive for both SIDT2 and Rab5(Q79L), exhibit key features of lysosomes.

**Fig. 3.**
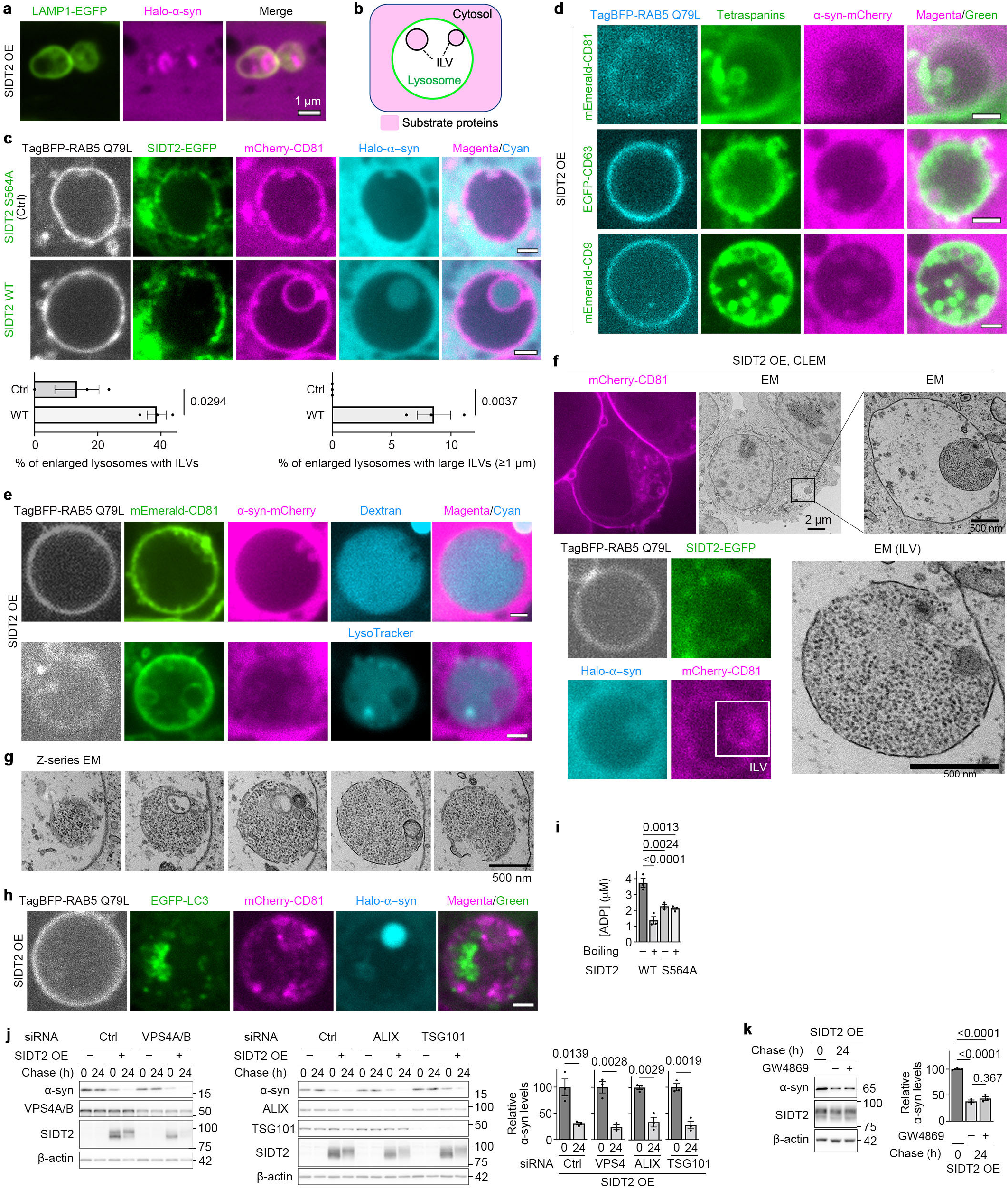
SIDT2 promotes microautophagic uptake of cytosolic proteins into lysosomes. (a) Localization of Halo-tagged α-synuclein within LAMP1–EGFP-positive lysosomes in SIDT2-overexpressing N2a cells. (b) Schematic of intraluminal vesicle (ILV) formation by membrane invagination (microautophagy). (c) ILV formation in enlarged endolysosomes in SIDT2-overexpressing cells (WT or S564A), visualised by mCherry-CD81 and internalized Halo-tagged α-synuclein. Percentage of enlarged lysosomes containing ILVs or large ILVs (≥1 µm). (d) α-synuclein–mCherry is detected inside ILVs labelled with fluorescently tagged CD81, CD63, or CD9 in Q79L RAB5-positive enlarged endolysosomes in SIDT2-OE cells. (e) Dextran or LysoTracker staining of enlarged endolysosomes; ILV structures are negative for both, indicating exclusion from the endolysosomal lumen. (f) Correlative light and electron microscopy (CLEM) images of enlarged endolysosomes in SIDT2-OE cells showing ILVs containing cytosolic components. (g) Z-series EM images showing multiple ILVs in enlarged endolysosomes of SIDT2-OE cells. (h) ILV membranes in enlarged endolysosomes of SIDT2-OE cells are negative for LC3. (i) ATPase activity of purified WT or SIDT2-S564A protein in vitro. (j, k) Effect of ESCRT component knockdown (VPS4A/B, ALIX, and TSG101) (j) or GW4869 treatment (k) on SIDT2-mediated α-synuclein degradation in N2a cells.

We next asked whether the SIDT2-dependent lysosomal protein uptake involves microautophagic membrane invagination. In cells co-expressing WT SIDT2, α-synuclein protein was detected within enlarged endolysosomes as punctate intraluminal structures surrounded by fluorescent protein-tagged tetraspanin protein CD81 (Fig. 3c), a marker of intraluminal membrane vesicles (ILVs)^22^. Importantly, compared with cells expressing the S564A mutant SIDT2, WT SIDT2 significantly increased the proportion of enlarged endolysosomes containing ILVs (ring-like structures; see Methods) (Fig. 3c), suggesting that SIDT2 promotes ILV formation through a Ser564-dependent activity. Notably, enlarged endolysosomes containing large ILVs (≥1 µm in diameter) were detected in WT SIDT2-expressing cells, whereas such structures were not observed in SIDT2(S564A)-expressing cells under the conditions tested (Fig. 3c). This suggests that SIDT2 contributes to the expansion of ILVs in an activity-dependent manner. Imaging analyses also showed that SIDT2 was largely retained on the membrane of enlarged endolysosomes and was rarely detected in ILVs (Fig. 3c), suggesting that SIDT2 may act at the limiting membrane to promote ILV formation rather than being incorporated into the resulting ILVs. To further examine whether internalized α-synuclein was associated with ILVs, we visualised ILVs with three different markers, fluorescent protein-tagged CD81, CD63, and CD9^22^. The internalized α-synuclein was surrounded by all three markers (Fig. 3d), indicating that the α-synuclein-containing structures are associated with ILV formation.

To assess whether these ILVs were generated by microautophagic invagination, we next examined whether they display topological features expected for such membrane invagination. First, using luminal staining of enlarged endolysosomes with dextran or LysoTracker, we found that the internal space of the large ILVs in cells expressing WT SIDT2 was negative for both dextran and LysoTracker, consistent with the sequestration of the cytosol by invagination of the endolysosomal membrane (Fig. 3e). Furthermore, correlative light and electron microscopy analysis revealed ILV membranes enclosing cytosolic components, including ribosomes, in SIDT2-overexpressing cells (Fig. 3f, g and Extended Data Fig. 8c; see also Movies S1 and S2). Together, these observations indicate that the large ILVs have a topology consistent with microautophagic sequestration of cytosol. In addition, an autophagosomal marker, LC3, was not observed on the ILV membrane (Fig. 3h), consistent with the idea that these ILVs do not derive from autophagosomal membranes. Altogether, these observations provide evidence that SIDT2 promotes microautophagic uptake of cytosolic components through the formation of enlarged ILVs.

Because Ser564 was required for SIDT2-mediated ILV formation, we next examined the catalytic activity associated with this conserved residue. Ser564 of SIDT2 has been predicted to constitute part of an evolutionarily conserved hydrolase-active site^19,20,23^. However, its endogenous hydrolytic substrate remains unknown. Using purified WT and SIDT2(S564A), we found that SIDT2 is capable of hydrolysing ATP, and Ser564 is critical for its ATPase activity (Fig. 3i). These findings raise the possibility that the ATPase activity of SIDT2 contributes to SIDT2-mediated microautophagy.

We next asked how this SIDT2-dependent pathway relates to previously described mechanisms of mammalian microautophagy and ILV formation. To date, knowledge of mammalian lysosomal microautophagy is limited. However, the endosomal sorting complex required for transport (ESCRT)-dependent lysosomal and endosomal microautophagy^24,25,26^ have been reported in mammalian cells. In addition, neutral sphingomyelinase (nSMase) has been implicated in ceramide-dependent ILV formation in endosomes^27^. Therefore, we investigated whether SIDT2-dependent enhancement of protein degradation requires ESCRT or nSMase. Knockdown of ESCRT proteins (VPS4, ALIX, and TSG101), pharmacological inhibition of ESCRT or nSMase by apilimod and GW4869, respectively, did not affect SIDT2-dependent enhancement of protein degradation in N2a cells (Fig. 3j, k and Extended Data Fig. 9a–c), suggesting that SIDT2-dependent microautophagic degradation can proceed independently of ESCRT and nSMase in N2a cells. To explore this further, we examined another cell line, SH-SY5Y, where overexpression of SIDT2 promoted protein degradation (Extended Data Fig. 9d). In SH-SY5Y cells, inhibition of ESCRT abolished SIDT2-dependent enhancement of protein degradation, whereas GW4869 treatment did not inhibit it (Extended Data Fig. 9e–g). These findings suggest that SIDT2-mediated microautophagic degradation is mechanistically distinct from nSMase-dependent ILV formation and exhibits cell-type-specific coupling to ESCRT machinery.

### Dominant-negative SIDT2 in neuromyopathy

Defects in intracellular degradation systems fundamental to homeostasis often lead to the development of diseases characterized by an accumulation of intracellular deposits, particularly in neurons and skeletal muscle^1^. We therefore investigated whether defects in SIDT2-mediated microautophagy are associated with protein aggregation diseases in humans. We searched our in-house database of whole-exome sequencing data from neuromyopathic patients and identified a heterozygous single-base deletion in *SIDT2* in one patient with familial neuropathy and distal myopathy (Fig. 4a). This patient was a 64-year-old Japanese man from a family showing dominant inheritance and was clinically diagnosed with neuropathy and distal myopathy with rimmed vacuoles (Supplementary text). Rimmed vacuolar myopathy (RVM)^28,29^ is a protein aggregation disease pathologically characterized by muscle fibres containing rimmed vacuoles, derived from cytosolic deposits which usually contain a variety of aggregation-prone proteins, including β-amyloid, ubiquitin, phosphorylated tau, and α-synuclein, and are surrounded by secondary accumulations of autophagosomes^30,31^. Computed tomography of the patient and histopathological analyses of a biopsy from the patient revealed typical features of RVM (Fig. 4b–f). Hamstring, triceps surae, and tibialis anterior muscles of the patient were preferentially replaced by fat (Fig. 4b). His vastus lateralis muscles showed marked variation in myofibre size, small group atrophy with several small angulated fibres and pyknotic nuclear clumps, fibrosis, and adipose tissue infiltration in the endomysium (Fig. 4c). Rimmed vacuoles, in which aggregation-prone proteins were characteristically accumulated along with autophagic and lysosomal proteins, were observed in some atrophic myofibres (Fig. 4d, e). Although skeletal muscle-specific conditional KO of *Sidt2* in mice has been reported to cause muscular dystrophy-like phenotypes, the patient’s blood creatine kinase levels were normal (186 IU/L). In addition to myogenic alterations, marked axonal loss in the intramuscular nerves was also observed (Fig. 4f). Whole-exome sequencing of gDNA revealed the presence of a heterozygous variant, c.2226delG in *SIDT2*. This mutation is predicted to cause a frameshift at Arg743, resulting in the replacement of 90 amino acid residues at the C-terminus with 72 amino acid residues of aberrant sequence (Extended Data Fig. 10a, b). The Sanger sequencing chromatogram showed comparable peak intensities for WT and mutant SIDT2 transcripts (Extended Data Fig. 10a). No mutations were identified in other genes known to cause RVM.

**Fig. 4.**
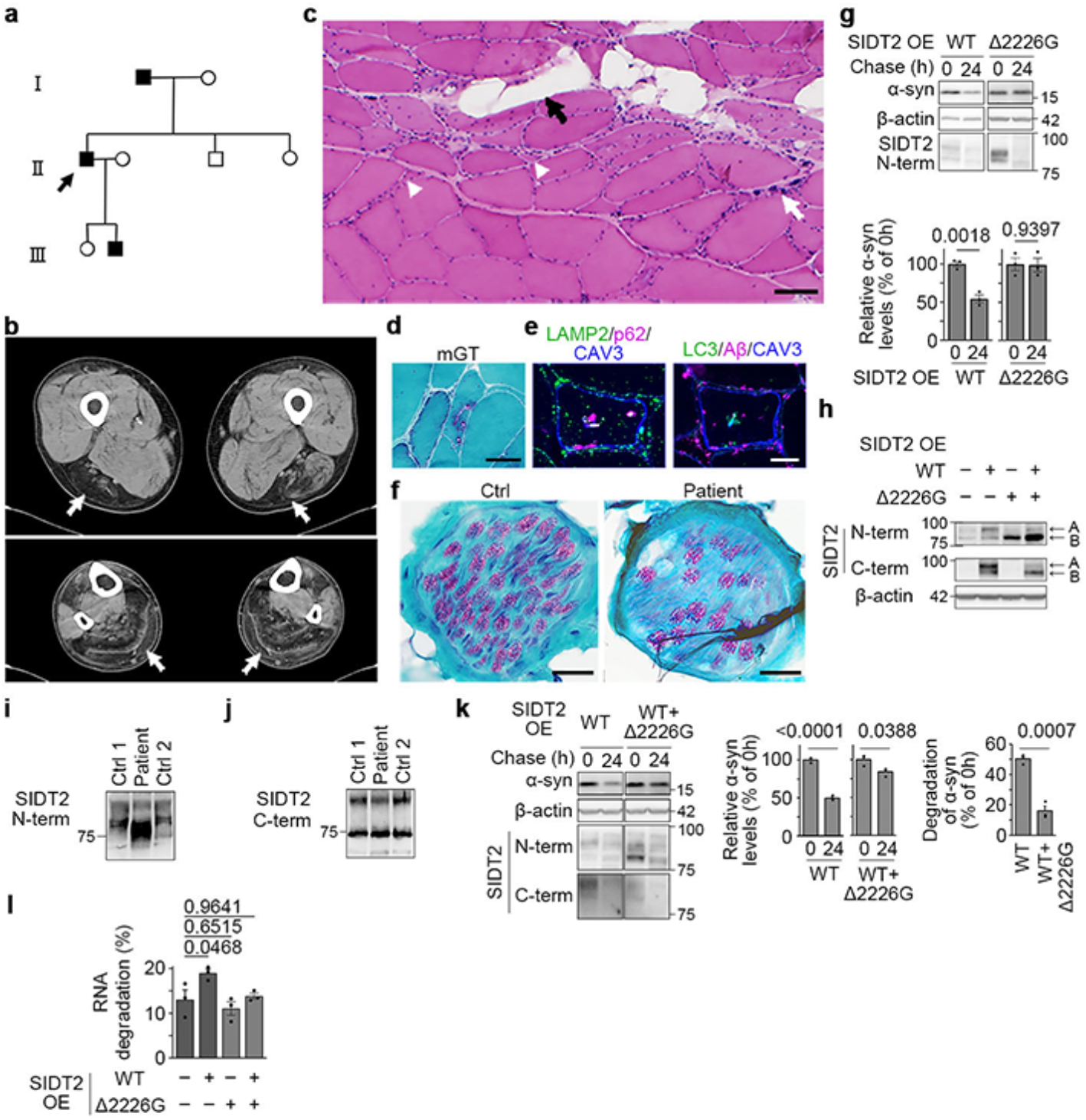
A dominant-negative SIDT2 mutation in rimmed vacuolar myopathy. (a) Pedigree chart of the proband and family affected by myopathy. The proband is indicated with an arrow. (b) Computed tomography (CT) image of the legs of the patient. The white arrow indicates the lesion. (c) Haematoxylin and eosin (HE)-stained section of patient-derived skeletal muscle. The white arrowheads indicate small angular fibres, the white arrow indicates pyknotic nuclear clumps, and the black arrow indicates adipose tissue infiltration. (d) Modified Gomori trichrome (mGT)-stained section of patient-derived skeletal muscle showing rimmed vacuoles. (e) Immunohistochemistry of skeletal muscle of the patient. (f) A mGT section of peripheral nerve in skeletal muscles of a control and patient biopsy. (g) Degradation of α-synuclein protein in cells overexpressing SIDT2 (WT or Δ2226G), using the Tet-Off system (n = 3). (h) Immunoblotting of N2a cell lysate overexpressing SIDT2 (WT, Δ2226G, or WT + Δ2226G), using specific antibodies against either the N- or C-terminus of SIDT2. (i, j) Immunoblotting of skeletal muscle of the patient using specific antibodies to either the N-terminus (i) or C-terminus (j) of SIDT2. (k) Degradation of α-synuclein protein in cells co-expressing SIDT2-WT and SIDT2-Δ2226G, compared with that of SIDT2-WT alone (n = 3). (l) Degradation of radiolabelled total RNA in cells, with overexpression as indicated (n = 3). Scale bars represent 40 µm.

To assess its pathogenicity, we first aimed to characterize the molecular properties of this mutant (Δ2226G) SIDT2. We observed that SIDT2-Δ2226G was incapable of increasing intracellular protein degradation (Fig. 4g), despite its normal localization to lysosomes (Extended Data Fig. 10c). SIDT2-Δ2226G also showed aberrant hypoglycosylation (Fig. 4h and Extended Data Fig. 10d). Immunoblotting of the patient muscle biopsy with an antibody recognizing both SIDT2-WT and SIDT2-Δ2226G showed decreased glycosylated SIDT2 (upper band) and increased hypoglycosylated SIDT2 (lower band) protein levels (Fig. 4i, j). However, overexpression of a SIDT2 mutant in which all four known N-glycosylation sites were substituted with glutamine (4NQ) promoted protein degradation to a similar extent as SIDT2-WT (Extended Data Fig. 10e), indicating that hypoglycosylation alone does not explain the reduction in SIDT2-dependent protein degradation by the Δ2226G mutant. Because the family history was consistent with autosomal dominant inheritance and SIDT2 forms homodimers, we speculated that SIDT2-Δ2226G may exert a dominant-negative effect. Notably, co-expression of SIDT2-Δ2226G severely impaired SIDT2-WT-induced protein and RNA degradation (Fig. 4k, l), indicating that SIDT2-Δ2226G is a dominant-negative mutation that impairs the SIDT2-dependent microautophagic degradation pathway. To test its pathogenic relevance in vivo, we next examined whether SIDT2 loss of function and the patient-associated mutation are sufficient to produce RVM-like pathology.

### RVM pathology across *Sidt2* mouse models

To examine the consequences of SIDT2 loss of function in vivo, we first generated *Sidt2*-deficient mice using the CRISPR/Cas9 system (Extended Data Fig. 11a). Importantly, the *Sidt2*-deficient mice recapitulated key features of RVM observed in the patient. The *Sidt2^−^*^/*–*^ mice exhibited reduced body size, locomotor activity, and grip strength (Fig. 5a and Extended Data Fig. 11b, c). A progressive and severe decline in muscle contraction and a gradual reduction in skeletal muscle size were also observed (Fig. 5b–e and Extended Data Fig. 11d). Histological analysis revealed myofibre atrophy and size variation, while no necrotic or regenerating fibres were observed (Fig. 5f), indicating that the phenotype is not dystrophic. In addition, fibrosis and adipocyte infiltration were also observed (Fig. 5f), resembling features seen in the patient muscle. Notably, rimmed vacuoles immunoreactive for various aggregation-prone proteins and LC3 were observed in the skeletal myofibres of *Sidt2^−^*^/*–*^ mice (Fig. 5g and Extended Data Fig. 11e). Biochemically, accumulation of α-synuclein was also observed in the skeletal muscle of these mice (Fig. 5h and Extended Data Fig. 11f). Ultrastructural analysis revealed amorphous inclusions surrounded by multilamellar bodies^30^ at the centre of myofibres in *Sidt2^−^*^/*–*^ mice (Fig. 5i). These findings indicate that *Sidt2*-deficient mice recapitulate key pathological features of RVM, rather than those of muscular dystrophy. Consistent with the accumulation of aggregation-prone proteins and amorphous inclusions, direct uptake and degradation of both protein and RNA were impaired in lysosomes derived from *Sidt2*-deficient muscles (Fig. 5j and Extended Data Fig. 12a). In line with this impairment, various RNAs were also accumulated in the *Sidt2*^−/–^ muscles (Extended Data Fig. 12b). These findings support the idea that impaired SIDT2-dependent microautophagic degradation underlies the RVM-like pathology in vivo. We also compared the histological and immunohistological features of muscles in skeletal muscle-specific conditional *Atg7*-KO mice with those of *Sidt2*-deficient muscle (Extended Data Fig. 12c, d). The *Sidt2*-deficient muscle showed more severe atrophy, fibrosis, and other myopathic features than *Atg7*-deficient muscle. Moreover, although the *Atg7*-deficient muscle also exhibited p62 accumulation, the accumulation pattern was diffuse and clearly distinct from that in *Sidt2*-deficient muscle where a small number of large deposits are often centrally localized within the muscle fibres. These observations further support that the myopathic features of *Sidt2*-deficient mice and the patient are unlikely to be the consequence of macroautophagy dysfunction.

**Fig. 5.**
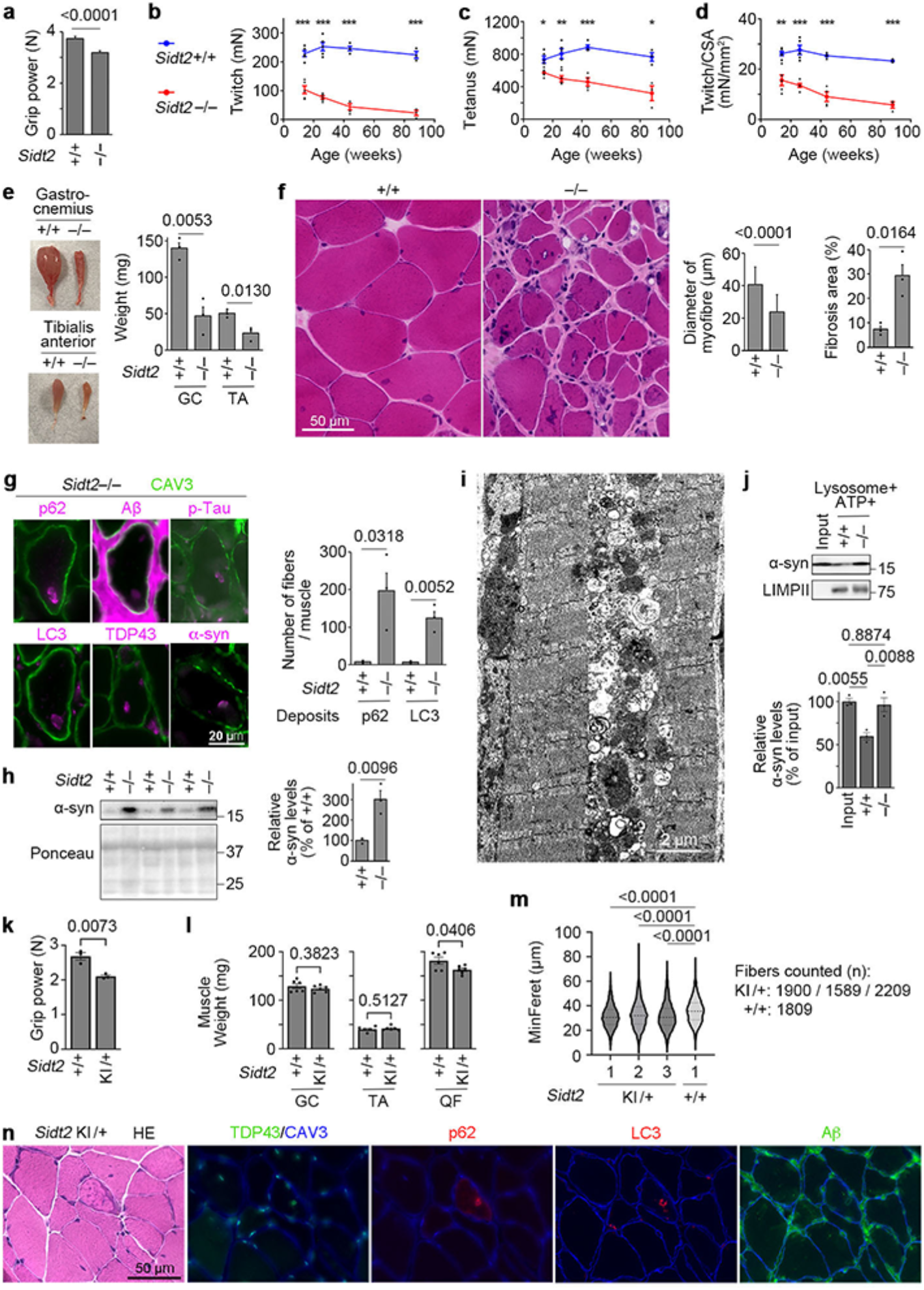
*Sidt2* deficiency impairs lysosomal microautophagic activity in skeletal muscle and promotes rimmed-vacuolar myopathic phenotype in mice. (a) Grip strength of *Sidt2*^+/+^ and *Sidt2^−^*^/*–*^ mice (+/+, n = 110 from 11 mice; –/–, n = 90 from 9 mice). (b–d) Changes in muscle contractions of gastrocnemius in *Sidt2*^+/+^ and *Sidt2^−^*^/*–*^ mice. Twitch (b), Tetanus (c), and Twitch/ cross-sectional area (CSA) (d) (n = 3–5). (e) Skeletal muscles of *Sidt2*^+/+^ and *Sidt2^−^*^/*–*^ mice and their weight (n = 3). \**P* < 0.05, \*\**P* < 0.001, \*\*\**P* < 0.0001. (f) HE sections of skeletal muscles of *Sidt2*^+/+^ and *Sidt2^−^*^/*–*^ mice and their quantification. Diameter of myofibres: +/+, n = 6,258; –/–, n = 5,350 from three mice. Fibrosis area, n = 3. (g) Immunohistochemistry of skeletal muscles of *Sidt2*^+/+^ and *Sidt2^−^*^/*–*^ mice (n = 3). (h) Immunoblotting of α-synuclein protein in skeletal muscles of *Sidt2*^+/+^ and *Sidt2^−^*^/–^ mice (n = 3). (i) Electron microscopy image of the ultrastructure of cytoplasmic inclusions in skeletal muscle of *Sidt2*^−/–^ mouse. (j) Degradation of α-synuclein protein by lysosomal preparations derived from skeletal muscles of WT or *Sidt2^−^*^/*–*^ mice (n = 3). (k) Grip strength of 98-week-old *Sidt2*^+/+^ and *Sidt2*^KI/+^ mice (n = 3). (l–n) Analysis of 104-week-old *Sidt2*^+/+^ and *Sidt2*^KI/+^ mice. Weights of muscles (n = 6) (l). Violin plot of Minimum Feret’s diameter (MinFeret) of myofibres in gastrocnemius; Comparison *P*-values appear above the plot, and horizontal lines identify the groups being compared. (m). Histological and immunofluorescence analysis of gastrocnemius from *Sidt2*^KI/+^ mice (n).

To directly test the pathogenicity of the patient-associated *SIDT2* mutation in vivo, we generated *SIDT2*-Δ2226G knock-in (KI) mice (Extended Data Fig. 13a–h). Notably, heterozygous KI mice exhibited reduced grip strength (Fig. 5k), decreased quadriceps femoris muscle weight (Fig. 5l), and muscle fibre atrophy (Fig. 5m). Again, we did not observe necrotic or regenerating fibres in these mice. Importantly, heterozygous KI mice exhibited accumulation of aggregation-prone proteins, surrounded by LC3, recapitulating the key features of rimmed vacuoles (Fig. 5n). We also observed myofibres positive for β-amyloid but negative for LC3, suggesting that amyloid deposition precedes accumulation of autophagosomes, as reported in other myopathies with rimmed vacuoles^30^ (Fig. 5n). In addition, homozygous KI mice showed significant reductions in body weight, grip strength, and muscle weights (Extended Data Fig. 13f–h). Together, these findings indicate that loss of *Sidt2* function is sufficient to produce RVM-like pathology in vivo and support a causal role for the patient-associated *SIDT2*-Δ2226G mutation.

## Discussion

In this study, we showed that SIDT2 promotes a lysosomal microautophagic pathway and delivers cytosolic proteins, including disease-associated aggregation-prone proteins, into lysosomes for degradation. We further identified a dominant-negative *SIDT2* mutation in familial neuromyopathy with rimmed vacuoles and showed that *Sidt2* deficiency and the patient-associated mutation recapitulate key pathological features of RVM in mice. At the cellular level, overexpression of SIDT2 markedly enhanced the degradation of cytosolic proteins, whereas SIDT2 knockdown reduced protein degradation. This SIDT2-dependent degradation pathway is lysosome-dependent; however, manipulation of SIDT2 did not alter luminal pH or proteolytic enzyme activity in lysosomes. The SIDT2-dependent degradation pathway was observed in both cultured cells and lysosomal fractions and was distinct from macroautophagy or CMA. Microscopic observations revealed that SIDT2 promotes ILV formation and enlargement, thereby facilitating the incorporation of cytoplasmic proteins into lysosomes for degradation.

SIDT2 has previously been predicted to belong to the hydrolase family^23^. Ser564 was identified as one of the putative catalytic residues, and recent cryo-electron microscopy studies structurally confirmed that Ser564 constitutes part of the catalytic centre^19,20^. In the present study, the S564A mutation impaired SIDT2-mediated protein degradation, and expression of the same mutant failed to induce enlargement of ILVs. Moreover, we showed in vitro that SIDT2 possesses ATPase activity, which was abolished in the S564A mutant. Because protein degradation by isolated lysosomes required ATP, likely reflecting a requirement for ATP during protein uptake, the ATPase activity of SIDT2 may contribute to membrane remodelling during microautophagic uptake. SIDT2 has also been reported to exhibit ceramidase and phospholipase activities in vitro^19,20^. How SIDT2 ATPase activity contributes to membrane remodelling during microautophagic invagination, and whether its reported lipid-hydrolytic activity is also involved, remain to be elucidated.

Although substrate proteins appeared as puncta within lysosomes upon SIDT2 overexpression (Fig. 3a), direct visualisation of microautophagic ILV formation in normal-sized lysosomes remains technically challenging because of the resolution limits of optical microscopy and rapid intraluminal degradation of ILVs. The Rab5(Q79L)-induced enlarged endolysosomes enabled visualisation of SIDT2-associated ILV formation. Advances in high-resolution imaging may enable direct visualisation of ILV formation in normal-sized lysosomes in the future.

SIDT2-dependent microautophagy showed a cell-type-specific dependency on the ESCRT machinery, the molecular basis of which remains to be determined. Recent studies, including our own, have reported both ESCRT-dependent^24,25^ and -independent^32^ forms of lysosomal microautophagy, suggesting that multiple mechanistically distinct microautophagy pathways may function in mammalian cells. In contrast, nSMase inhibition did not suppress SIDT2-dependent degradation in either cell type, suggesting that this pathway is distinct from canonical ceramide-dependent ILV formation. An ESCRT-dependent endosomal pathway has also been reported to deliver ubiquitinated α-synuclein to lysosomes^33,34^, suggesting that multiple mechanisms contribute to the maintenance of proteostasis.

Previously, SIDT2 was shown to mediate the lysosomal uptake of RNA and DNA through processes termed RNautophagy and DNautophagy, respectively^5,10^. Given that SIDT2 promotes a lysosomal microautophagic pathway, RNautophagy and DNautophagy may represent nucleic acid forms of SIDT2-dependent microautophagy. This idea is consistent with recent cryo-electron microscopy studies, revealing that SIDT2 does not adopt channel-like structures^19,20^. Moreover, the SIDT2-dependent protein degradation pathway described here shares key features with RNautophagy and DNautophagy, including the requirement for Ser564 and ATP. This similarity further supports the idea that nucleic acid uptake mediated by SIDT2 also occurs via microautophagy.

SIDT2 has been implicated in a late stage of macroautophagy, particularly in autophagosome–lysosome fusion, although the underlying mechanism remains unclear^7,8^. Thus, the molecular role of SIDT2 in macroautophagy has not been established. In contrast, our study shows that SIDT2 regulates lysosomal microautophagy. Furthermore, our data suggest that macroautophagy is regulated in a compensatory manner upon perturbation of SIDT2-dependent microautophagy. One possible explanation for the discrepancy with previous reports is that the changes in macroautophagy observed upon SIDT2 perturbation may not reflect a direct regulatory role of SIDT2, but rather a secondary response to impaired proteostasis.

The secondary accumulation of autophagosomes is a known feature of RVMs, which arise from cytosolic protein accumulation rather than primary macroautophagic defects. Consistently, macroautophagy-deficient muscles did not harbour rimmed vacuoles, exhibiting pathological features distinct from those of *Sidt2*-deficient muscles (Extended Data Fig. 12d). This is also the case for muscles deficient in CMA (muscle-specific *Lamp2a* KO), which did not harbour rimmed vacuoles but instead exhibited distinct pathological features, including mitochondrial abnormalities and tubular aggregates^35^.

RVM pathology is generally attributed to cytoplasmic accumulation of proteins and other substances that should constitutively be degraded; abnormalities in the luminal functions of lysosomes, such as luminal pH and hydrolase activity, are not considered the primary cause of this group of myopathies^28^. This is consistent with the fact that causative genes of RVMs reported previously encode extra-lysosomal proteins. To our knowledge, SIDT2 is the first gene causally linked to RVM that encodes a lysosomal protein. By contrast, known causative genes for autophagic vacuolar myopathies, a distinct group of myopathies, encode lysosomal proteins, and are thought to be primarily caused by lysosomal aberrations^28^. Importantly, none of these primary lysosomal myopathies are characterized by rimmed vacuoles. The fact that both *Sidt2* deficiency and the dominant negative *SIDT2*-Δ2226G mutation cause the formation of rimmed vacuoles strongly suggests that SIDT2 dysfunction leads to RVM through impaired microautophagic uptake of degradation substrates from the cytoplasm into the lysosomal lumen (Extended Data Fig. 14), rather than through primary defects in lysosomal luminal acidification or proteolytic activity.

SIDT2-dependent microautophagy may be part of an integrated system for cellular proteostasis. Such systems are thought to be particularly important for the quality control of cellular proteins in postmitotic long-lived tissues during stress conditions, including aging and disease. Therefore, the microautophagic pathway identified in this study may provide a potential therapeutic target for degenerative diseases in muscles and neurons.

## Methods

### siRNAs

The following siRNAs were purchased from JBioS with targeting sequences against EGFP (siCtrl: 5′-CAGCACGACUUCUUCAAGU-3′), mouse SIDT2 (siSIDT2: 5′-GAGUUUCCGUCCAGUAUUU-3′), mouse SIDT2 (siSIDT2-2: 5′-GUACCUCUACCAAAAAGUG-3′), mouse LAMP2A (5′-GACTGCAGTGCAGATGAAG-3′), mouse VPS4A (5′-CAACCCUAGCGUUAUGAUU-3′), and mouse VPS4B (5′-CUACCUUGCGAGUGCUACA-3′). The following siRNAs were purchased from Sigma-Aldrich (Merck) with targeting sequences against mouse ALIX (5′-GGAUUACUUUGGCGAUGCU-3′), mouse TSG101 (5′-GGUACAAUCCCAGUGCGUU-3′), human VPS4A (5′-CCGAGAAGCUGAAGGAUUA-3′), and human VPS4B (5′-CCAAAGAAGCACUGAAAGA-3′).

### Plasmids

pCI-neo vectors containing EGFP, and WT and mutant forms of mouse SIDT2 were prepared as described previously^5,9^. pEGFP–LC3, pEGFP–LAMP1, and pCI-neo–human CHMP2B–Q165X, which expresses the dominant-negative form of CHMP2B were described previously^36–38^. cDNA encoding human *SIDT2* was subcloned into the pCI-neo mammalian expression vector (Promega, E1841). pCI-neo–human SIDT2–Δ2226G was generated via mutagenesis using Pfu Turbo DNA Polymerase (Agilent, 600250), in accordance with the manufacturer’s instructions. pTRE–Tight vectors harbouring each of the FLAG-tagged reporter proteins, pTRE–Tight–α-synuclein–mCherry, pTRE–Tight–α-synuclein–EGFP–mCherry, and pmCherry–N1–human SIDT2–Δ2226G, were generated by subcloning each construct into pTRE–Tight or pmCherry–N1 vector, respectively. pTRE–Tight–GAPDH–ΔNR was generated by introducing a mutation into pTRE–Tight–GAPDH by mutagenesis. pCAG-neo–His_8_–GFP–mSIDT2 was generated by inserting a synthetic DNA fragment encoding His_8_ and GFP tags after amino acid 69 of mouse SIDT2 in a pCI-neo-based construct, which was subsequently subcloned into the pCAG–Neo vector (Fujifilm Wako, 163-25601). pCI-neo–mouse 4NQ– SIDT2 was generated by replacing the WT mouse SIDT2 coding sequence with a synthetic DNA sequence containing four point mutations (N54Q, N60Q, N123Q, and N165Q), synthesised by Integrated DNA Technologies (IDT). The pPAmCherry-KFEFQ plasmid was constructed by inserting a DNA sequence encoding 21 amino acids of ribonuclease A (MKETAAAKFERQHMDSSTSAA) into the pPAmCherry–N1 Vector (Takara, Z2584N). pTagBFP–human RAB5B-Q79L was generated by subcloning the cDNA encoding human RAB5B into the pTagBFP-N vector (Evrogen, FP172), followed by introduction of the Q79L mutation via mutagenesis. All resulting plasmid constructs were analysed by sequencing. Plasmids encoding fluorescently tagged tetraspanin proteins, including pmEmerald–CD9 (Addgene, 54029), pEGFP–CD63 (Addgene, 62964), pmEmerald–CD81 (Addgene, 54031), and pmCherry–CD81 (Addgene, 55012), were used to label intraluminal vesicles (ILVs). pFN21A–Halo–α-synuclein (Promega, FHC02635) was used to express human α-synuclein fused to an N-terminal HaloTag.

### Cell lines

The *Atg5*^−/–^, *Atg13*^−/–^, and *Atg5*^+/+^ mouse embryonic fibroblasts (MEFs) were provided by Professor N. Mizushima (The University of Tokyo)^39,40^. HeLa Tet-Off Advanced cells were purchased from Takara (Clontech, 631156). 293FT cells were purchased from Invitrogen (Thermo Fisher Scientific, R70007). C2C12 cells were purchased from ATCC. Neuro2a (N2a) cells were provided by Dr. Y. Suzuki^36^. *Atg13*- and *Atg5*-KO N2a cells described previously^38^ were used. *LAMP2*-KO HeLa cells described previously^5^ were also used. N2a–tau cells, which stably express human full-length 2N4R tau (P301L) and GFP–K18 (Q244–E372, P301L), and harbour tau aggregates introduced by K18 tau fibrils (P301L), were prepared as described^41^. *Sidt2*^−/–^ and *Sidt2*^+/+^ MEFs were generated using mouse embryos of the respective genotype on embryonic days 13.5–14.5 and immortalized by transfection of SV40 large T antigen-expressing vector (pBSSVD2005, Addgene, 21826). All cell lines were grown in Dulbecco’s modified Eagle’s medium (DMEM, Gibco, C11995500) supplemented with 10% foetal bovine serum. All cell lines used in this study were confirmed to be mycoplasma-free using the MycoStrip Mycoplasma Detection Kit (InvivoGen).

### Antibodies

The following primary antibodies were used in this study: mouse monoclonal anti-DYKDDDDK tag (FLAG) antibody (1E6, Wako, 012-22384), mouse monoclonal anti-β-actin antibody (AC-15, Sigma-Aldrich, A1978), rabbit monoclonal anti-ATG5 antibody (EPR1755(2), Abcam, ab108327), rabbit polyclonal anti-ATG13 antibody (Sigma-Aldrich, SAB4200100), rabbit monoclonal anti-LC3B antibody (D11, Cell Signaling Technology, 3868S), rabbit polyclonal anti-α-synuclein antibody (Sigma-Aldrich, AB5038P), mouse monoclonal anti-Tau antibody (Tau46, Cell Signaling Technology, 4019S), mouse monoclonal anti-phospho-Tau antibody (AT8, Absolute Antibody, Ab02331-23.0), rabbit polyclonal anti-LAMP2A antibody (Abcam, ab18528), mouse monoclonal anti-VPS4A antibody (VPS4-110, Sigma-Aldrich, SAB4200215), rabbit monoclonal anti-LIMPII antibody (EPR12080, Abcam, ab176317), and rabbit polyclonal anti-SIDT2 antibody raised against the N-terminus of human SIDT2 (Abnova, PAB27211). The production of rabbit polyclonal antibody against the N-terminus of mouse SIDT2 (NVSQKDAEFERTYA) was outsourced to Eurofins. Rabbit polyclonal antibody against the C-terminal region of human and mouse SIDT2 (DLDTVQRDKIYVF) was produced as previously described^9^.

### Immunoblotting

Protein samples were separated by SDS-PAGE and transferred onto PVDF membranes (Bio-Rad, 1620177; or Wako, 033-23433), which were then blocked with 3% bovine serum albumin prepared in Dulbecco’s PBS or 5% skim milk prepared in PBS containing 0.1% Tween 20. This was followed by incubation with primary antibodies prepared in either of the blocking buffers overnight at 4 °C. The membranes were then washed with 0.1% Tween 20 prepared in PBS and probed with horseradish peroxidase-conjugated secondary antibodies (Thermo Fisher Scientific [Pierce], 31430 or 31460). Signals were then visualised using ImmunoStar Zeta (Wako, 295-72404) or ImmunoStar LD (Wako, 290-69904) and detected using the FluorChem chemiluminescence imaging system (Alpha Innotech) or FUSION chemiluminescence imaging system (Vilber-Lourmat). The signal intensity was quantified using FluorChem or FUSION software.

### Measuring degradation levels of radiolabelled total proteins in cultured cells

A total of 3.0 × 10^4^ cells/well of *Atg5^+/+^* or *Atg13*^−/–^ MEFs, or 2.0 × 10^4^ cells/well of *Atg5^−/–^* MEFs, were seeded in 24-well plates (FALCON, 353047). The following day, 37 kBq/well leucine, L-[4,5-^3^H(N)]-(PerkinElmer, NET135H) was added to the medium and the cells were radiolabelled for 48 h. After 48 h of labelling, the cells were washed with 500 µL/well of medium containing 8 mM unlabelled leucine (Sigma-Aldrich) and incubated in another 500 µL/well containing the same medium for the indicated period (16 or 24 h). Following this incubation, the cells were harvested using 500 µL/well of trypsin (Gibco, 25200-056) and mixed with the same volume of 10% trichloroacetic acid (TCA, Wako, 207-04955). The samples were then chilled on ice, centrifuged at 6,500×*g* and 4 °C for 10 min, and the TCA-insoluble fraction was collected. The same procedures were also performed after the labelling (0 h), and TCA-insoluble radioactivity for all samples was measured using Tri-Carb 3100TR (Packard). The remaining intracellular protein was calculated relative to the TCA-insoluble radioactivity at 0 h and expressed as a percentage. The degradation levels of protein were measured as the relative decline in radioactivity between 0 and 24 h. For the knockdown experiments, 10 pmol/well siRNA was transfected into cells 4 h prior to radiolabelling using Lipofectamine 3000 (Thermo Fisher Scientific, L3000-015), in accordance with the manufacturer’s instructions.

### Measuring protein degradation in cultured cells using Tet-Off system

A total of 1.5 × 10^5^ cells/well of N2a cells (WT, *Atg5*-KO, or *Atg13*-KO) or HeLa Tet-Off Advanced cells were seeded in 12-well plates (FALCON, 353043). The cells were transfected with plasmids using polyethyleneimine (Polysciences, 24765-2) the following day, and 1 µg/mL doxycycline (Clontech, 631311) was added 24 h later to stop the expression of the FLAG-tagged reporter proteins. The cells were then incubated for 8 h to exclude the effect of mRNA degradation, and the samples defined as “0 h” in the experiment were collected. The cells for samples at other time points were further incubated and collected at each of the indicated time points. The relative levels of FLAG-tagged reporter proteins were quantified by immunoblotting using an anti-FLAG antibody, and the degradation levels were calculated by subtracting the relative levels of proteins at each time point (percent in relation to the 0 h time point) from 100%. N2a cells were transfected with 0.16 µg/well of pTet-Off vector, 0.02 µg/well of pTRE–Tight vector expressing each FLAG-tagged reporter protein, and pCI-neo vector expressing EGFP or mouse SIDT2 (WT, 3YS, S564A, or 4NQ). For experiments using cycloheximide, 50 µg/mL cycloheximide (Sigma-Aldrich, C1988-1G) was added to the medium at 0 h. For the experiment shown in Fig. 4g, N2a cells were transfected with 0.10 µg/well of pTet-Off vector, 0.02 µg/well of pTRE–Tight-α-synuclein-FLAG, and 0.02 µg/well of pCI-neo-SIDT2-WT or 0.08 µg/well of pCI-neo–SIDT2– Δ2226G. For the experiment shown in Fig. 4k, N2a cells were transfected with 0.10 µg/well of pTet-Off vector, 0.02 µg/well of pTRE–Tight–α-synuclein–FLAG, 0.02 µg/well of pCI-neo–human SIDT2–WT, and 0.08 µg/well of pCI-neo vector expressing EGFP or human SIDT2–Δ2226G. The 4:1 ratio of Δ2226G to WT plasmid was chosen to achieve approximately equal protein expression levels of the two constructs, consistent with the comparable SIDT2 protein levels observed between the patient and control muscle biopsies (Figs. 4i, j). For the knockdown experiments, N2a cells were transfected with 0.10 µg/well of pTet-Off vector, 0.05 µg/well of pTRE–Tight-α-synuclein-FLAG, 0.05 µg/well of pCI-neo vector expressing EGFP or mouse SIDT2, and 10 pmol/well of siRNA using Lipofectamine 3000, in accordance with the manufacturer’s instructions. HeLa Tet-Off Advanced cells were transfected with 0.10 µg/well of pTRE–Tight–α-synuclein-FLAG and 0.10 µg/well of pCI-neo vector expressing EGFP or mouse SIDT2.

### Proteoliposome reconstitution

For affinity-based isolation of FLAG-SIDT2, immunoprecipitation was performed as described by Contu et al.^9^, unless otherwise indicated. Cells were harvested 24 h post-transfection and lysed on ice in 1% (v/v) C12E9 (CAS: 3055-97-8) in base buffer (50 mM HEPES–KOH, pH 7.6, 150 mM KCl, 5% [v/v] glycerol) containing protease inhibitors (Complete, EDTA free; Roche Applied Science, 1873580). Lysates were clarified by centrifugation, and the supernatant was incubated with anti-FLAG M2 agarose affinity gel (Sigma-Aldrich) for 1 h at 4 °C with rotation. Beads were washed twice with lysis buffer and once with elution buffer (0.1% [v/v] C12E9 in base buffer). Proteins were eluted using 100 µg/mL FLAG peptide (Wako) in elution buffer for 30 min at 4 °C with rotation. Eluates were dialyzed against the same buffer for 24 h at 4 °C (1:1000 sample-to-buffer ratio), with one buffer change after 12 h, using Slide-A-Lyzer Dialysis Cassettes (Thermo Fisher).

Proteoliposomes were prepared as previously described^42^, with some modifications. Briefly, *Escherichia coli* polar lipid extract (Avanti, 100600C) and 18:1 (Δ9-Cis) PC (DOPC) (Avanti, 850375C) were mixed at a 7:3 ratio (weight/weight) with a total concentration of 20 mg/mL in chloroform. The chloroform was then evaporated using nitrogen gas, and the remaining lipids were suspended in ice-cold reconstitution buffer (50 mM HEPES, 150 mM KCl, 5 % glycerol, pH 7.6), followed by sonication using a Handy Sonic (TOMY, UR-21P) to produce small unilamellar vesicles, which were subjected to freezing and thawing for three cycles to generate large multilamellar vesicles. The large multilamellar vesicles were then filtered through polycarbonate Membrane 0.1 µm (Avanti, 610005) for 11 rounds using Extruder Set (Avanti, 610000), and large unilamellar vesicles (LUVs) were collected. Then the reconstitution buffer was added to dilute the LUVs to a concentration equivalent to 4 mg/mL of original lipid, and C12E9 was mixed into the solution at 0.3% of final concentration to destabilize the LUVs. The optimal concentration of C12E9 was determined by measuring the turbidity of the solution. The destabilized LUVs were then mixed with FLAG-SIDT2 protein at 100:1 ratio (weight/weight) of lipid and protein, and the mixed solution was rotated at 4 °C overnight. The following day, 40 mg/mL of Bio-Beads SM-2 (Bio-Rad, 152-3920) was added to the solution three times, every hour, and the mixed solution was then rotated at 4 °C overnight to remove the detergent. The beads were removed from the solution, and the resulting proteoliposomes were pelleted by ultracentrifugation at 4 °C for 30 min at 267,000×*g.* The proteoliposomes were resuspended in the reconstitution buffer at a concentration equivalent to 4 mg/mL of original lipid and subjected to subsequent experiments.

### Fluorescence microscopy

For the experiment shown in Fig. 1l, 3.0 × 10^5^ cells/dish of N2a cells were seeded in 35 mm glass-bottomed dishes (IWAKI, 3970-035), transfected with 0.32 µg/dish of pTet-Off vector, 0.04 µg/dish of pTRE–Tight–α-synuclein–mCherry, and 0.04 µg/dish of pCI-neo (empty vector) or pCI-neo–mouse SIDT2, and 1 µg/mL doxycycline was added after 1 d. The following day, the cells were treated with LysoTracker Green DND-26 (Life Technologies, L7526) to label the lysosomes and then analysed using an FV10 confocal microscope (Olympus [Evident]). Quantification and calculation of colocalization rates were performed with ImageJ software via the JACoP plugin. For the experiment shown in Extended Data Fig. 3a, 3.0 × 10^5^ cells/dish of N2a cells were seeded in 35 mm glass-bottomed dishes. The following day, the cells were transfected with 0.28 µg/dish of pTet-Off vector, 0.04 µg/dish of pTRE–Tight-α-synuclein-mCherry, 0.04 µg/dish of pEGFP-N1-LAMP1, and 0.04 µg/dish of pCI-neo-mouse SIDT2 and then treated with 1 µg/mL doxycycline 1 d thereafter. The following day, the fluorescence in single cells was analysed using DeltaVision Elite (Applied Precision [Cytiva]). For the experiment shown in Extended Data Fig. 3b, 3.0 × 10^5^ cells/dish of N2a cells were seeded in 35 mm glass-bottomed dishes. One day later, the cells were transfected with 0.32 µg/dish of pTet-Off vector, 0.04 µg/dish of pTRE–Tight–α-synuclein–EGFP–mCherry, 0.04 µg/dish of pCI-neo (empty vector), or pCI-neo–mouse SIDT2 and then treated with 1 µg/mL doxycycline the day after to stop the expression of α-synuclein–EGFP–mCherry. The next day, the cells were treated with LysoTracker BlueDND-22 (Life Technologies, L7525) to label the lysosomes and then analysed using an FV10 confocal microscope. For the experiment shown in Extended Data Fig. 7a, 3.0 × 10^5^ cells/dish of N2a cells were seeded in 35 mm glass-bottomed dishes. The following day, the cells were transfected with 0.04 µg/dish of PAmCherry–KFERQ, 0.04 µg/dish of LAMP1–EGFP, and 0.32 µg/dish of pCI-neo (empty vector). The next day, PAmCherry–KFERQ was photoactivated by UV illumination for 5 min, and the cells were then incubated either in starvation medium (amino acid-free medium [Wako 048-33575] without serum) or in nutrient-rich (complete) medium for 24 h and analysed using an FV1000 confocal microscope (Olympus [Evident]). For the experiment shown in Extended Data Fig. 7b, 3.0 × 10^5^ cells/dish of N2a cells were seeded in 35 mm glass-bottomed dishes. The following day, the cells were transfected with 0.04 µg/dish of PAmCherry–KFERQ, 0.04 µg/dish of LAMP1– EGFP, and either 0.32 µg/dish of pCI-neo (empty vector) or 0.04 µg/dish of pCI-neo–SIDT2 together with 0.28 µg/dish of pCI-neo. The next day, PAmCherry–KFERQ was photoactivated by UV illumination for 5 min, and the cells were then incubated in complete medium for 24 h and analysed using an FV1000 confocal microscope. For the experiment shown in Extended Data Fig. 10c, 3.0 × 10^5^ cells/dish of N2a cells were seeded in 35 mm glass-bottomed dishes, and the cells were transfected with 0.16 µg/dish of pmCherry–N1–human SIDT2–Δ2226G with 0.24 µg/well of pCI-neo (empty vector). The following day, the cells were treated with LysoTracker Green DND-26 to label the lysosomes and then analysed using an FV10 confocal microscope.

### Measurement of autophagic flux

For the measurement of the autophagic flux at steady-state levels shown in Fig. 1o, 1.5 × 10^5^ cells/well of WT N2a cells were seeded in 12-well plates. The following day, the cells were transfected (using polyethyleneimine) with 0.16 µg/well of pCI-neo (empty vector), 0.02 µg/well of pCI-neo (empty vector), or pCI-neo–mouse *Sidt2*, 0.02 µg/well of pMRX–IP–GFP–LC3–RFP–LC3ΔG^43^, and then the fluorescence was measured two days later using a SpectraMax i3x (Molecular Devices). Autophagic flux was measured as the ratio of fluorescence between GFP and RFP in each well. To analyse whether the cells are capable of driving autophagy^40^ in Extended Data Fig. 1a or 4b, the protocol was as follows. Briefly, MEFs (WT, *Atg5*-KO, or *Atg13*-KO) and N2a cells (WT or *Atg13*-KO) were seeded in 12-well plates, and the medium was replaced with starvation medium with or without 100 nM bafilomycin A1, or normal medium as a control, for 2 h. The samples were then subjected to immunoblotting against LC3, and the autophagic flux was assessed based on the accumulation of LC3-II in the presence of bafilomycin A1.

### Imaging of enlarged endolysosomes and ILVs

N2a cells were seeded onto 35-mm glass-bottomed dishes (ibidi) and transfected using polyethyleneimine with the indicated plasmids. Cells were imaged 20–24 h after transfection using a super-resolution microscope (spinSR10, Olympus [Evident]). For Fig. 3a, cells were transfected with 0.04 μg of Halo-α-synuclein, 0.04 µg of LAMP1–EGFP, and 0.04 μg of SIDT2–WT. For Fig. 3c, cells were transfected with 0.04 μg of Halo-α-synuclein, 0.04 μg of mCherry–CD81, 0.08 μg of Q79L RAB5, and 0.04 μg of either SIDT2–WT or SIDT2– S564A, together with 0.04 µg of SIDT2(WT) –EGFP or SIDT2(S564A) –EGFP, respectively. For Fig. 3d, cells were transfected with 0.04 μg of α-synuclein–mCherry, 0.04 μg of SIDT2–WT, 0.08 μg of BFP–RAB5 Q79L, and 0.04 μg of mEmerald–CD81, EGFP–CD63, or mEmerald–CD9. For Fig. 3e, cells expressing SIDT2–WT, BFP–RAB5 Q79L, mEmerald–CD81, and α-synuclein–mCherry were incubated with Alexa Fluor 647-labeled dextran for 16 h (final 0.1 mg/mL) (Thermo Fisher Scientific, D22914) or LysoTracker Green DND-26 (Thermo Fisher Scientific, L7526) for 30 min before imaging. For Fig. 3h, cells were transfected with 0.04 μg of EGFP–LC3, 0.04 μg of mCherry–CD81, 0.08 μg of Q79L RAB5, 0.04 μg of Halo– α-synuclein, and 0.04 μg of WT SIDT2. For visualisation of Halo-tagged proteins, cells were incubated with 5 μM HaloTag ligand TMR (Promega, G8251) or 1 μM HaloTag SaraFluor 650T Ligand (Goryo chemical, A308-01), for 15 min at 37 °C, followed by washing with fresh medium before imaging. All imaging was performed under live cell conditions, unless otherwise noted. For ILV quantification, ILVs were defined as mCherry–CD81-positive ring-like structures. Structures showing diffuse mCherry–CD81 fluorescence within the lumen, rather than a ring-like pattern, were excluded from analysis. Each data point represents the percentage of enlarged endolysosomes containing ILVs calculated from ten cells, with three measurements obtained from two independent dishes per condition.

### Correlative light and electron microscopy

N2a cells were seeded onto 35-mm glass-bottomed dishes (MatTek, P35G-1.5-14-C/H) and transfected using polyethyleneimine with 0.04 μg of Halo–α-synuclein, 0.04 μg of mCherry–CD81, 0.08 μg of BFP–RAB5 Q79L, 0.04 μg of SIDT2(WT) –EGFP, and 0.04 μg of SIDT2–WT. Cells were imaged 20–24 h after transfection using a super-resolution microscope (spinSR10). To identify cells of interest, markings were made by scraping part of the cell layer, followed by capturing Nomarski images. The cells were then fixed by a mixture of 2% formaldehyde and 2.5% glutaraldehyde in 0.1 M HEPES–NaOH buffer (pH 7.4) containing 1 mM CaCl_2_ for more than 2 h. Fixed samples were extensively rinsed with HEPES–NaOH buffer with 1 mM CaCl_2_, followed by rinsing with 0.1 M sodium cacodylate buffer (pH 7.4) containing 1 mM CaCl_2_. Post-fixation was carried out with 1% osmium tetroxide and 0.1% potassium ferrocyanide in 0.1 M sodium cacodylate buffer with 1 mM CaCl_2_ for 1 h. Samples were washed with distilled water, dehydrated through a graded ethanol series, and infiltrated with Epon812 resin. Polymerization was performed at 60 °C for at least 48 h. Serial ultrathin sections were prepared using a Leica EM UC7 ultramicrotome and mounted onto Formvar-coated copper grids. After counterstaining with lead citrate, sections were imaged using a JEM-1400Flash transmission electron microscope.

### Lysosomal uptake/degradation of protein

Experiments using lysosomal preparations were performed as previously described^5,6,9,10,12,44,45^ with slight modifications. Briefly, lysosomal preparations were obtained from MEFs, N2a cells, mouse brain, or skeletal muscles using the Lysosome Enrichment Kit for Tissue and Cultured Cells (Thermo Scientific, 89839) and suspended in 300 mM sucrose. Lysosomes were then mixed with 0.1 µg/assay of recombinant α-synuclein (BIOMOL, SE-256), tau (tau-441, Wako, 205-20331), or SOD1 protein and 10 mM 3-(N-morpholino)propanesulphonic acid (Dojin, 345-01804) buffer (pH 7) with or without 10 mM (final concentration) adenosine 5′-triphosphate disodium salt n-hydrate (Wako 017-09673) in 300 mM sucrose, and incubated at 37 °C for 0, 5, 10, or 20 min. The samples were then subjected to immunoblotting, and the relative levels of α-synuclein, tau, or SOD1 were quantified. The degradation of α-synuclein was obtained by subtracting the relative levels of α-synuclein (percent of input) from 100%. For the experiment shown in Fig. 2d, the suspended lysosomes were preincubated with or without 50 µg/mL pepstatin A (Peptide Institute, 4397-v) and 50 µg/mL E64d (Peptide Institute, 4321-v) for 30 min on ice and then subjected to the following experiments. For the experiments shown in Fig. 2g, the suspended lysosomes were preincubated with or without 1.0 µg/assay of recombinant His-tagged Hsc70 protein for 30 min on ice and then subjected to the following experiments. For the experiments shown in Fig. 2h, lysosomes were mixed with 0.1 µg of recombinant α-synuclein, together with 5 µg of total RNA or dNTPs. For the experiment shown in Extended Data Fig. 5d, lysosomes were incubated at 37 °C for 20 min in the presence or absence of ATP and precipitated by centrifugation (4 °C, 17,700×*g*, 1 min). The supernatant was collected and then incubated with 0.1 µg/assay of recombinant α-synuclein protein at 37 °C for 20 min. For the experiment shown in Extended Data Fig. 6c, lysosomes suspended in 300 mM sucrose were dissolved in citrate buffer containing 1% Triton X (final volume), and then incubated with 0.1 µg/assay of recombinant α-synuclein at 37 °C for 5 min. For the experiments shown in Fig. 5j and Extended Data Fig. 12a, lysosomes were prepared from skeletal muscles of the lower extremities of mice and subjected to the following experiments. For RNA degradation, 5.0 µg/assay of total RNA was used. In the experiments using lysosomes derived from distinct samples, the lysosome amount was confirmed by immunoblotting with LIMPII.

### Measurement of lysosomal pH

A total of 2.0 × 10^5^ cells/well of *Sidt2*^+/+^ and *Sidt2^−^*^/*–*^ MEFs were seeded in 35 mm glass-bottomed dishes, and cells were treated with or without 100 nM bafilomycin A1, followed by 15 min of treatment with 1 µM LysoSensor Green DND-189 (Thermo Fisher Scientific, L7535) 1 h later. The medium was then discarded, cells were incubated for 1 h, and lysosomal pH was analysed as relative fluorescence units using an FV10 confocal microscope and the ImageJ software.

### Immunoelectron microscopy

Immunoelectron microscopy was performed as previously described^5,6,10,12^. Briefly, lysosomal preparations were incubated in the presence of ATP with or without recombinant α-synuclein protein and 5 nm gold particles (Cytodiagnostics, G-5-20) and then precipitated by centrifugation. Lysosomes were then fixed with 0.1% glutaraldehyde and 4% paraformaldehyde in 0.1 M phosphate buffer overnight at 4 °C, dehydrated in a series of water/ethanol mixtures to 100% ethanol, and subsequently embedded in LR White (Nisshin EM, 3962). The samples were then sectioned at a thickness of 100 nm and collected on 400-mesh nickel grids. The sections were subsequently immunolabelled with anti-α-synuclein antibody (Cell Signaling Technology, 2628 for Fig. 2e and Extended Data Fig. 5f; and Abcam, ab138501 for Extended Data Fig. 5e, g), followed by 15 nm gold-labelled goat anti-rabbit IgG (H+L) (Amersham), and observed using a Tecnai Spirit transmission electron microscope (FEI [Thermo Fisher Scientific]) at 80 kV.

### Preparation ΔDQ mutant α-synuclein

The ΔDQ α-synuclein recombinant protein was prepared as previously described^6^, with slight modifications. Briefly, Rosetta2 (DE3) competent cells (Millipore, 71397) were transformed with pGEX-6P-1-ΔDQ α-synuclein, and protein expression was induced with 500 µM isopropyl-β-D-1-thiogalactopyranoside (Wako, 094-05144). The cells were incubated at 25 °C overnight and harvested by centrifugation. The cells were then suspended in PBS containing 1% Triton X and protease inhibitors and lysed by sonication. The sample was then centrifuged for 10 min at 18,000×*g* and 4 °C, and GST-tagged ΔDQ α-synuclein was purified from the supernatant using Glutathione Sepharose 4B (GE Healthcare [Cytiva], 17-0756-01). ΔDQ α-synuclein protein was then cleaved from GST using PreScission Protease (GE Healthcare [Cytiva], 27-0843-01).

### ATPase activity assay

His8-GFP-SIDT2 protein (WT or S564A) was transiently expressed in 293FT cells. Cells were harvested 24 h after transfection. Frozen pellets of large amounts of harvested cells (60 dishes of φ150 mm dish) were homogenized in a high-osmolarity buffer (10 mM HEPES pH 7.5, 30 mM NaCl) and sonicated to disrupt the cells. After ultracentrifugation (200,000×*g*, 4 °C, 30 min), the pellets were further disaggregated by homogenization and sonication and then subjected to a second ultracentrifugation. The pellets and the membrane protein fraction were suspended in 10 mM HEPES, 100 mM NaCl (pH 7.5) containing 10% glycerol. The membrane protein fractions were solubilized by incubation in solubilization buffer (final concentration: 1% n-dodecyl-β-D-maltoside and 0.1% cholesteryl hemisuccinate, 50 mM HEPES, 150 mM NaCl; pH 7.5) containing protease inhibitor with gentle rotation for 60 min at 4 °C. The first purification step of the WT and S564A SIDT2 proteins was performed using a column packed with TALON Metal affinity resin (Clontech, 635503), taking advantage of the His-tag fused to SIDT2. Each sample was eluted with an elution buffer (200 mM imidazole, 50 mM Tris-HCl, 0.1 M NaCl) containing 0.03% n-dodecyl-β-D-maltoside and 0.0003% cholesteryl hemisuccinate. The eluted fraction was then concentrated using an Amicon Ultra-15 tube (Merck). In the second purification step, the crude SIDT2 protein fraction was further purified by fluorescence-detection size-exclusion chromatography, utilizing the fluorescence of GFP-fused SIDT2, which was run on an HPLC system (Shimadzu, Prominence LC-20 system) connected to a Superose-6 Increase column (Cytiva, 29-0915-96). Fractions corresponding to the major peak (elution time: 35–45 min) of GFP fluorescence intensity (Ex 480 nm, Em 530 nm) were detected using the LabSolutions system (Shimadzu), and the fractions were collected and concentrated using VIVASPIN500 (Sartorius, Z614025). After concentration, the solution was aliquoted and stored at –80 °C.

The ATPase activity of SIDT2 (WT or S564A) was measured using the malachite-based BIOMOL Green reagent (Enzo Life Sciences, BML-AK111). The purified proteins (0.7 µg/sample) were incubated at 37 °C for 60 min in 50 µL of reaction buffer containing 1 mM ATP, 20 mM HEPES (pH 7.5), 2 mM MgCl_2_, 150 mM NaCl, and 0.03% DDM. After incubation, the reaction was terminated by adding 100 µL of BIOMOL Green reagent, then incubated for 20 min to develop the colour. The inorganic phosphate (Pi) released was measured by absorbance at 620 nm using a SpectraMax i3x microplate reader. A phosphate standard curve was prepared according to the manufacturer’s protocol and used to calculate the Pi concentration. To confirm that the activity depended on the native structure of SIDT2, protein samples were boiled (100 °C, 5 min) and then cooled before the assay. ATPase activity of WT and S564A SIDT2 was compared under native and heat-denatured conditions.

### Ethical considerations

All clinical information and materials used in this study were obtained for diagnostic purposes with written informed consent. The ethics committee of the National Center of Neurology and Psychiatry (NCNP) approved this study and the use of human subjects for this study (approval number B2024-131). All relevant ethical regulations were followed. We reviewed the medical information obtained from interviews conducted by physicians and evaluated the patient’s clinical manifestations, disease course, and laboratory data.

### Skeletal muscle imaging

Skeletal muscle magnetic resonance imaging data in the Digital Imaging and Communications in Medicine format were available as part of the clinical information at the time of muscle pathology diagnosis at the NCNP.

### Mutation analysis

At the NCNP, whole-exome sequencing analysis has been performed in 2,131 undiagnosed cases of suspected hereditary neuromuscular disease using genomic DNA from blood or biopsied muscle samples^46^. We screened this whole-exome sequencing database for the variants in SIDT2 and evaluated them according to the following variables: (1) mutation effect (i.e., splicing, start lost, stop gained or lost, frameshift, nonsynonymous codon change, and codon insertion or deletion); and (2) variation frequency, < 0.001 in a public database [Genome Aggregation Database (gnomAD), 1000 Genomes Project, or Exome Sequencing Project v. 6500 (ESP6500)], and <0.01 for in-house data^47^. The identified variants were confirmed by Sanger sequencing. Expression of the transcript with variants was analysed by direct sequencing and cloning of the RT-PCR products of the SIDT2 transcript from the patient-derived skeletal muscles. Primer sequences for the PCR are available upon request.

### Experimental animals

All animal care and experimental procedures were reviewed and approved by the Animal Care and Use Committee of the National Institute of Neuroscience (approval numbers: 2026025 and 2026010). All experiments were performed in accordance with the Guidelines for Animal Experiments of the National Institute of Neuroscience and with the relevant national regulations, including the Act on Welfare and Management of Animals in Japan. The study was conducted in compliance with the ARRIVE guidelines. Mice were maintained under specific pathogen-free conditions in a controlled environment (22 ± 2 °C; 12-h light/dark cycle; ad libitum access to food and water). For starvation, mice were placed in clean cages without food for 30 h (from 9 AM to 3 PM the next day). Water was provided ad libitum. Starvation for 30 h resulted in a 10% loss of weight, but did not result in any mortalities.

*Atg7*-flox mice were developed by Prof Komatsu Masaaki (Juntendo University)^48^ and MuCre-A mice were developed by Prof Kyoko Koishi (University of Otago)^49^ (20877696). Both mouse models were provided by the Riken BioResource Research Center (RBRC02759 and RBRC01386). Mice with skeletal muscle-specific *Atg7* KO were generated by mating.

*Sidt2* knockout and *Sidt*2 Δ2226G knock-in mice were generated in this study. B6C3F1 mice used for fertilized egg collection and ICR females used as pseudopregnant recipients were purchased from SLC Japan. C57BL/6J mice used for backcrossing were purchased from CLEA Japan. One-cell-stage zygotes were obtained by mating B6C3F1 stud males with superovulated females. Donor females were deeply anesthetized with a mixture of medetomidine, midazolam, and butorphanol and euthanized by cervical dislocation before oviduct collection. Pronuclear microinjection and mouse transgenesis were performed using standard procedures. Recipient females were anaesthetized with the same anaesthetic regimen for oviductal embryo transfer, and anaesthesia was reversed with atipamezole to allow recovery.

For lysosome isolation from the brain and skeletal muscle in in vitro reconstitution experiments, as well as for skeletal muscle weight measurements, mice were deeply anesthetized with sevoflurane, and tissues were collected following exsanguination. For histological analyses, immunostaining, electron microscopy, and RNA extraction, mice were deeply anaesthetized with a mixture of medetomidine, midazolam, and butorphanol, and skeletal muscles were collected following exsanguination.

### Muscle pathology

Muscle samples were collected from the deltoid muscle of the patient and from the quadriceps femoris, gastrocnemius, and tibialis anterior muscles of mice and then frozen in isopentane cooled in liquid nitrogen^50^. Histological analysis was performed as previously described^51^. Electron microscopy-based observation of the hamstring muscles from the mice was performed in line with a previously reported protocol^52^. Immunofluorescence of the skeletal muscles was performed in accordance with previously reported methods^30^. For the morphometric analysis, we stained frozen transverse sections of gastrocnemius muscles for caveolin 3, LC3, and p62 and acquired images under a Keyence BZ X-700 microscope. Five randomly selected images (per mouse) with caveolin 3 staining were used to measure fibre diameters using ImageJ (NIH). The minimal inner diameters of 2,000 myofibres from each mouse were measured. Images of whole muscles were used to quantify LC3- and p62-positive deposits^53^. MinFeret values were measured using ImageJ on cross-sectional muscle images. The following primary antibodies were used in this study: anti-LAMP2 mouse-Mono (H4B4, Santa Cruz), anti-p62 Rb-poly (BML-PW9860, Enzo), anti-LC3 Rb-poly (NB100-2220, Novus), anti-Cav3 goat-poly (N-18 Santacruz), anti-amyloid mouse-Mono (6E-10, Covance), and anti-TDP-43 Rb-poly (Protein Tech).

### De-glycosylation of SIDT2 using PNGase F

N2a cells expressing GFP or human SIDT2 (WT, Δ2226G, or WT + Δ2226G) were lysed with 1% Triton X-100 lysis buffer (50 mM Tris-HCl, pH 7.5, 150 mM NaCl, 5 mM EDTA, and 1% Triton X-100) containing protease inhibitors, and treated with PNGase F (New England BioLabs, P0704S) at 37 °C for 4 h or left untreated. Following the treatment, the samples were subjected to immunoblotting.

### Generation of *Sidt2* knockout mice

*Sidt2^−^*^/*–*^ mice were generated using the CRISPR/Cas9 system. The sequences for sgRNA against two distinct sites in exons 13 (5′-ACTGGACTCCATGAGCTCCG-3′) and 26 (5′-CGACTTGGACACAGTACAGC-3′) of SIDT2 were designed using the web-based CRISPR design tool CRISPOR (https://crispor.gi.ucsc.edu/) and cloned into the pX330 plasmid (Addgene, 42230). The plasmids were injected into the pronuclei of B6C3F1 fertilized eggs. The genotype of the resulting KO mice (deletion between exons 13 and 26) was confirmed by PCR with the following primers: 5′-GGGACCTCTCCTACAGTTACCAGG-3′, 5′-CCGCAGAGGCAACTCCAAGGCTC-3′, and 5′-CGCAGAGTCCAGGGAGCACAAC-3′. The KO mice were backcrossed with C57BL/6J mice for at least eight generations before being used in the experiments; the data shown in Fig. 5e–i and Extended Data Fig. 12a, b, d are exceptions as in these cases, backcrossing was done for three generations. We confirmed that rimmed vacuoles were also observed in the backcrossed mice (eight generations). Phenotypic analyses were performed at 81–100 weeks of age unless otherwise noted. All phenotypic analyses were performed using female mice.

### Generation of *Sidt2* <u>Δ</u>2226G KI allele

To generate KI mice carrying the human patient’s c.2226delG mutation (a frameshift at Arg743), two guide RNAs, depicted in Extended Data Fig. 13a, were designed to delete exons 24 to 26 of the mouse *Sidt2* gene and simultaneously KI the human patient-derived frameshifted exon 24–26 sequence. For the design of Guide RNAs 1 and 2, the CRISPOR tool (https://crispor.gi.ucsc.edu/) was employed to select guide sequences closest to the deletion sites with minimum off-target effects. Two parts of the CRISPR guide RNAs, crRNA and tracrRNA, were chemically synthesised (Integrated DNA Technologies). The synthesised RNA sequences were as follows. Guide RNA 1-crRNA: 5’-GCUUGAUCCUCUCGCCGCUCguuuuagagcuaugcu-3’, Guide RNA 2-crRNA: 5’-AUGUCUUCUAGCAGCAUCUGguuuuagagcuaugcu-3’, and tracrRNA: 5’-AGCAUAGCAAGUUAAAAUAAGGCUAGUCCGUUAUCAACUUGAAAAAGUGGCACCGAGUCGGUGCUUU-3’. For pronuclear injection, equimolar crRNA and tracrRNA were mixed and annealed. Recombinant Cas9 protein (Alt-R S.p. HiFi Cas9 Nuclease V3) was purchased from Integrated DNA Technologies. The targeting vector containing the human patient-derived frameshifted Exon 24-26 sequence flanked by 1 kb homology arms on both sides was artificially synthesised by Eurofins Genomics.

The mixture of annealed crRNA/tracrRNA for Guide RNA1 and Guide RNA 2, Cas9 protein, and the targeting vector was injected into mouse fertilized eggs at the concentrations of 25, 25, 100, and 15 ng/μL, and the eggs were transferred into the oviducts of surrogate mothers to raise pups.

Genomic DNAs were prepared from newborns’ tails by proteinase K treatment in lysis buffer (Takara Bio). KI mice were screened by PCR using Tks Gflex DNA polymerase (Takara Bio). To exclude the possibility that the donor DNA had been randomly integrated into the genome, a subset of genotyping primers was designed external to the donor DNA’s homology arms. For KI candidates, the PCR products were analysed by Sanger sequencing to confirm that they carried the correct insertions. The obtained founders were backcrossed with B6C3F1 to get F1 heterozygous progenies, which were again sequence-verified to harbour the designed KI allele. For genotyping PCR, Primer F (5’-TGTTGGTTCAGGGATGTGGGTATATGG-3’) and Primer R (5’-CCACCAGGAACTATCCTGAGTCAGG-3’) are used. A 2,498 bp product was amplified from the WT *Sidt2* allele, whereas a 574 bp product was amplified from the KI allele. F1 heterozygous KI mice derived from two independent founder lines were used for analysis. Age-matched B6C3F1 mice were used as WT controls.

Homozygous KI mice were generated either by crossing two independent KI founder mice or by intercrossing F1 heterozygous KI mice derived from different founder lines. For analysis, mice derived from two distinct founder pairs (with different male and female founders) were used. WT controls for homozygous KI mice were littermates obtained from F1 × F1 intercrosses. All phenotypic analyses were performed using female mice.

### Physiological examination of mice

Grip strength of the combined forelimbs or hindlimbs and voluntary locomotion within an individual cage using a running wheel were measured as described previously^51^. The contractile properties of the tibialis anterior and gastrocnemius muscles were measured in accordance with previously reported protocols^54,55^.

### Expression in *Sidt2*^+/+^ and *Sidt2^−^*^/*–*^ mice

RNA was purified from frozen sections of mouse gastrocnemius using the TRI Reagent (Sigma-Aldrich, T9424), in accordance with the manufacturer’s instructions. cDNA was then synthesised using PrimeScript RT Reagent Kit with gDNA Eraser (Perfect Real Time) (TaKaRa, RR047) and employed as a template for quantitative PCR on the CFX96 Touch Real-Time PCR Detection System (Bio-Rad). TB Green *Premix Ex Taq* II (Tli RNaseH Plus) (TaKaRa, RR820) was used for performing the quantitative PCR. The levels of each target RNA were normalized to those of *Gapdh*. The following primers were used for the detection of respective RNAs: 28S rRNA Forward: 5′-TGGGAATGCAGCCCAAAGC-3′, Reverse: 5′-CCTTACGGTACTTGTTGACTATCG-3′; 18S rRNA Forward: 5′-GTAACCCGTTGAACCCCATT-3′, Reverse: 5′-CCATCCAATCGGTAGTAGCG-3′; *App* Forward: 5′-GGAAGCAGCCAATGAGAGAC-3′, Reverse: 5′-GGTTCATGCGCTCGTAGATCA-3′; *p62* Forward: 5′-TGTGGAACATGGAGGGAAGAG-3′, Reverse: 5′-GTGCCTGTGCTGGAACTTTC-3′; α-synuclein Forward: 5′-ATGGATGTGTTCATGAAAGGAC-3′, Reverse: 5′-AACCACTCCTTCCTTAGTTTTG-3′; *Tau* Forward: 5′-CCTGAGCAAAGTGACCTCCAAG-3′, Reverse: 5′-CAAGGAGCCAATCTTCGACTGG-3′; *Tardbp* (TDP-43) Forward: 5′-CTCAGTGTATGAGAGGAGTCCGAC-3′, Reverse: 5′-ACACTATGAGGTCAGATGTTTTCTGG-3′; and *Map1lc3b* (LC3) Forward: 5′-GACGGCTTCCTGTACATGGTTT-3′, Reverse: 5′-TGGAGTCTTACACAGCCATTGC-3′. Specific primers for 5S rRNA, 5.8S rRNA, and *Gapdh* were prepared as previously described^5^.

### Statistical analyses

Quantitative data were analysed using the unpaired Student’s *t*-test for comparison between two groups and Dunnett’s or Tukey’s multiple comparison tests for more than two groups. For count data (immunoelectron microscopy gold particle numbers), the Mann–Whitney U test was used. All statistical tests were two-sided. For most experiments, each data point represents a measurement from a biologically independent sample. For image-based quantifications, individual cells, lysosomes, or muscle fibres from the same biological sample were quantified as separate data points. For immunoblotting and for microscopy-based images with quantitative data, representative images are shown. Bar graphs display mean values with error bars representing s.e.m., unless otherwise specified. Data points are shown unless omitted for clarity due to large sample size. Violin plots were generated using GraphPad Prism to visualise the data distribution. Each plot shows a smoothed density of the data, with the thick central dotted line indicating the median and the thinner dotted lines indicating the first and third quartiles.

## Acknowledgments

We thank N. Mizushima for constructive advice and suggestions that improved our work, and for providing *Atg5*^+/+^, *Atg5^−^*^/–^, and *Atg13^−/–^* MEFs and pMRX-IP-GFP-LC3-RFP-LC3ΔG, M. Komatsu for providing *Atg7*-flox mice, K Koishi for providing MuCre-A mice, and H. Osakada, M. Fujiwara, K. Hase, M. Shikama, and Y Hara for technical assistance. We also thank Editage and Edanz Group for editing the English text of a draft of this manuscript.

## Author contributions

Conceptualization: Y.F., S.N., T.K.; Formal analysis: Y.F., V.R.C., C.K., M.O., S.N., T.K.; Funding acquisition: Y.F., V.R.C., I.K.-H., G.M., N.N.N., I.N., K.W., T.F., S.N., T.K.; Investigation: Y.F., V.R.C., C.K., M.O., H.F., H.K., R.S., M.S., I.K.-H., M.I., Y.O., Y.I.-U., T.I.; Methodology: Y.F., V.R.C., R.S., I.K.-H., G.M., K.M., N.N.N., K.W., T.F., S.N., T.K.; Project administration: Y.F., S.N., T.K.; Supervision: I.N., S.N., T.K.; Validation: Y.F., V.R.C., C.K., S.N., T.K.; Visualisation: Y.F., M.O., S.N., T.K.; Writing - original draft: Y.F. and S.N., T.K.; and Writing - review and editing: Y.F., S.N., T.K.

## Data and material availability

All data generated and used in this study are either included in this article or are available from the corresponding authors upon reasonable request. Correspondence and requests for materials should be addressed to T.K. or S.N.

## Supplementary information

### Supplementary text

#### Clinical Presentation

The patient’s symptoms began in the third decade of life with gradual progression of muscle weakness and atrophy in the upper and lower extremities. The patient developed an inability to run at the age of 54 years owing to a bilateral foot drop, and an inability to perform fine handiworks at 59 years of age. The patient’s father and son were also affected (Fig. 4a). The genetic evidence in this study is based on a single patient, and co-segregation analysis in other affected family members was not possible, because the affected relatives did not provide consent for genetic analysis. Electromyography and nerve conduction revealed early recruitment and markedly lower amplitudes of motor unit action potential and compound muscle action potential.

Upon radiography of the patient, the hamstring, triceps surae, and tibialis anterior muscles were observed to be preferentially replaced by fat (Fig. 4b). His vastus lateralis muscles showed marked variation in myofibre size, small group atrophy with several small angulated fibres and pyknotic nuclear clumps, fibrosis, and adipose tissue infiltration in the endomysium (Fig. 4c). Rimmed vacuoles, in which aggregation-prone proteins were characteristically accumulated along with autophagic and lysosomal proteins, were observed in some atrophic myofibres (Fig. 4d, e). In addition to those myogenic observations, a remarkable deficit in axons of the intramuscular nerves was also observed (Fig. 4f).

---

**Extended Data Fig. 1.**
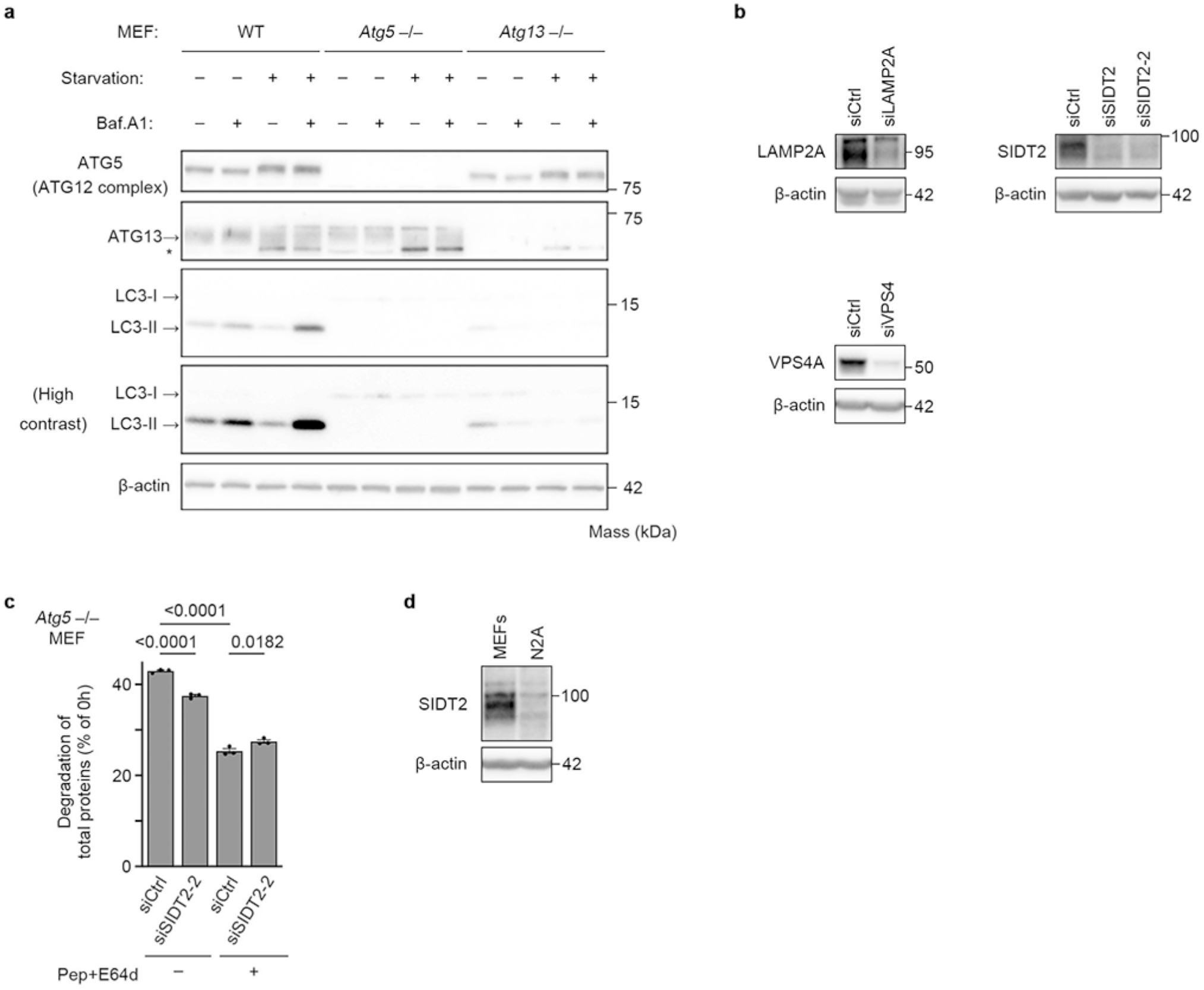
SIDT2 mediates lysosomal uptake and degradation of proteins in cells. (a) Immunoblotting of the same MEF lines showing the absence of ATG5 or ATG13, and LC3 flux under nutrient-rich or starvation conditions with or without bafilomycin A1 (100 nM, during 2-h starvation), confirming impairment of macroautophagy in both knockout lines. *, nonspecific bands. (b) Expression levels of LAMP2A, SIDT2, and VPS4A in *Atg5^−/–^* MEFs knocked down for respective gene products. (c) Effect of SIDT2 knockdown and lysosomal protease inhibitors on degradation of radiolabelled total proteins in *Atg5^−/–^* MEFs, using another siRNA against SIDT2 (siSIDT2-2) (n = 3). (d) Comparison of SIDT2 protein expression between MEFs and N2a cells by immunoblotting.

**Extended Data Fig. 2.**
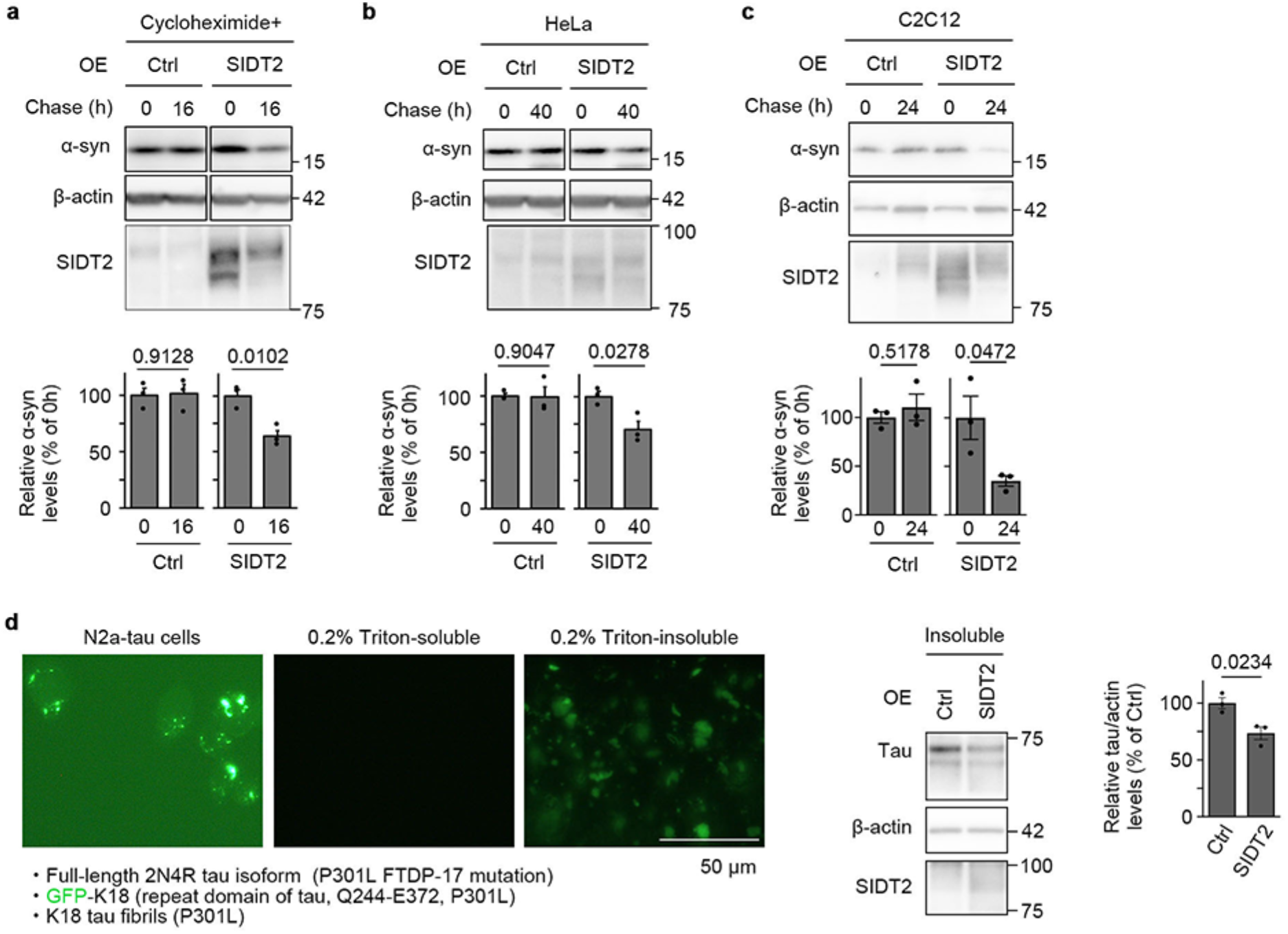
SIDT2 mediates lysosomal degradation of proteins. (a) Degradation of α-synuclein protein in N2a cells overexpressing SIDT2 using the Tet-Off system, under cycloheximide treatment to additionally inhibit protein synthesis during the chase (n = 3). (b, c) Degradation of α-synuclein protein in HeLa (b) or C2C12 (c) cells overexpressing SIDT2, using the Tet-Off system (n = 3). (d) Effect of SIDT2 overexpression on tau aggregates. Fluorescence microscopy images of GFP–tau in N2a cells harbouring tau aggregates (N2a–tau) before and after extraction with lysis buffer (50 mM Tris [pH 7.5], 1 mM EDTA, 150 mM NaCl, 0.2% Triton X-100) containing protease inhibitors (left). Aggregated tau signals were exclusively detected in the Triton-insoluble fraction and were not detected in the Triton-soluble fraction. Immunoblot analysis of Triton-insoluble fractions from control or SIDT2-OE cells shows reduced levels of insoluble tau upon SIDT2 expression (right), suggesting that SIDT2 promotes the clearance of tau aggregates, either directly or indirectly (e.g., via degradation of soluble tau). Quantification of tau levels normalised to β-actin is shown (n = 3).

**Extended Data Fig. 3.**
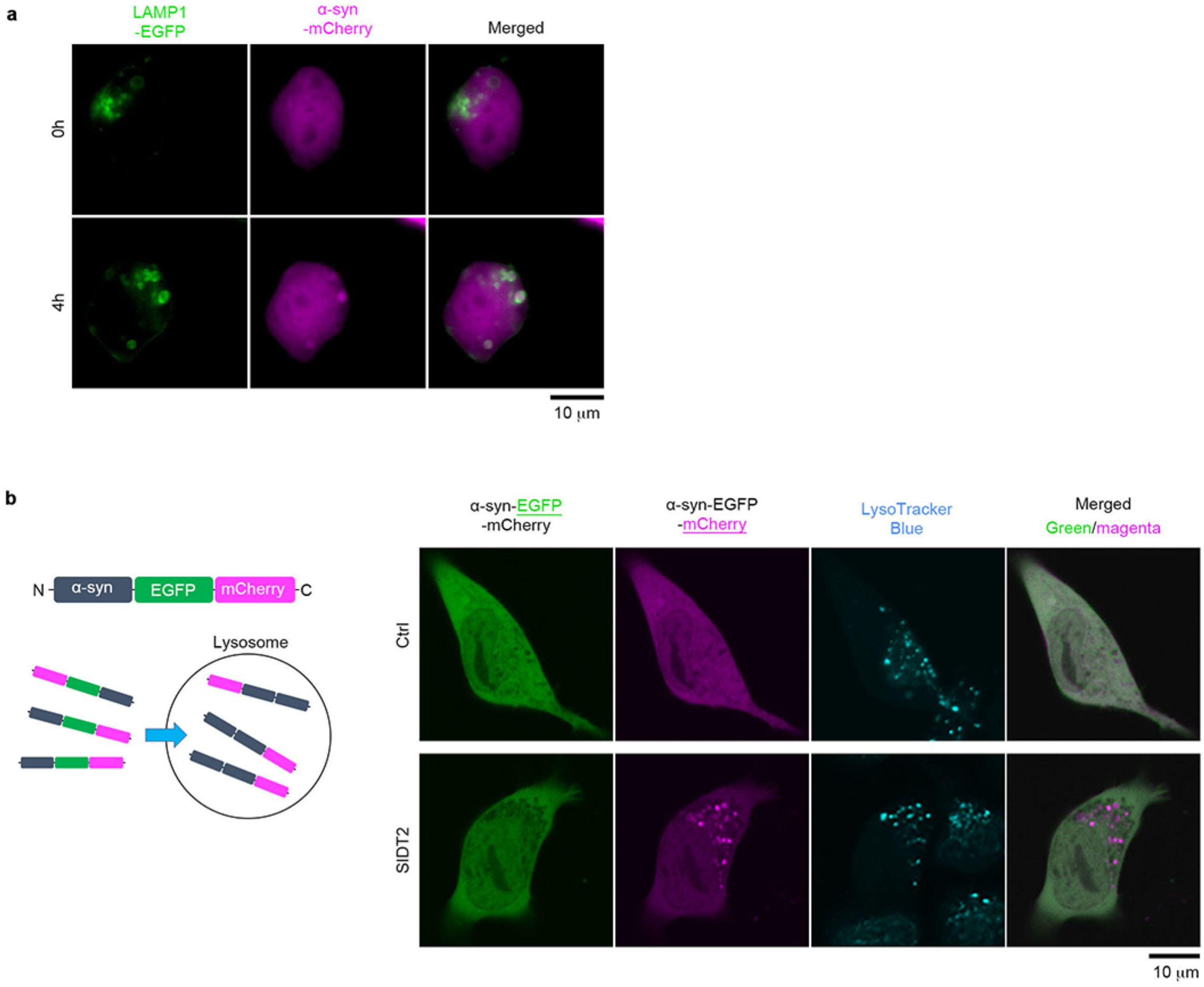
Localization of fluorescent protein-labelled α-synuclein protein in SIDT2-overexpressing cells. (a) Time-lapse imaging of α-synuclein–mCherry in an N2a cell overexpressing SIDT2. α-synuclein–mCherry is primarily distributed in a diffuse state at 0 h and subsequently appears in lysosomes at 4 h. (b) Left panel: Schematic of the α-synuclein–EGFP–mCherry construct. When α-synuclein–EGFP–mCherry is delivered into acidic organelles such as lysosomes, the EGFP signal is selectively quenched, leaving behind mCherry-positive only puncta prior to degradation of the constructs. Right panel: Localization of α-synuclein–EGFP–mCherry protein in SIDT2-overexpressing N2a cells. EGFP-negative but mCherry-positive puncta that localize to lysosomes indicate that the protein is imported into the lumen of lysosomes.

**Extended Data Fig. 4.**
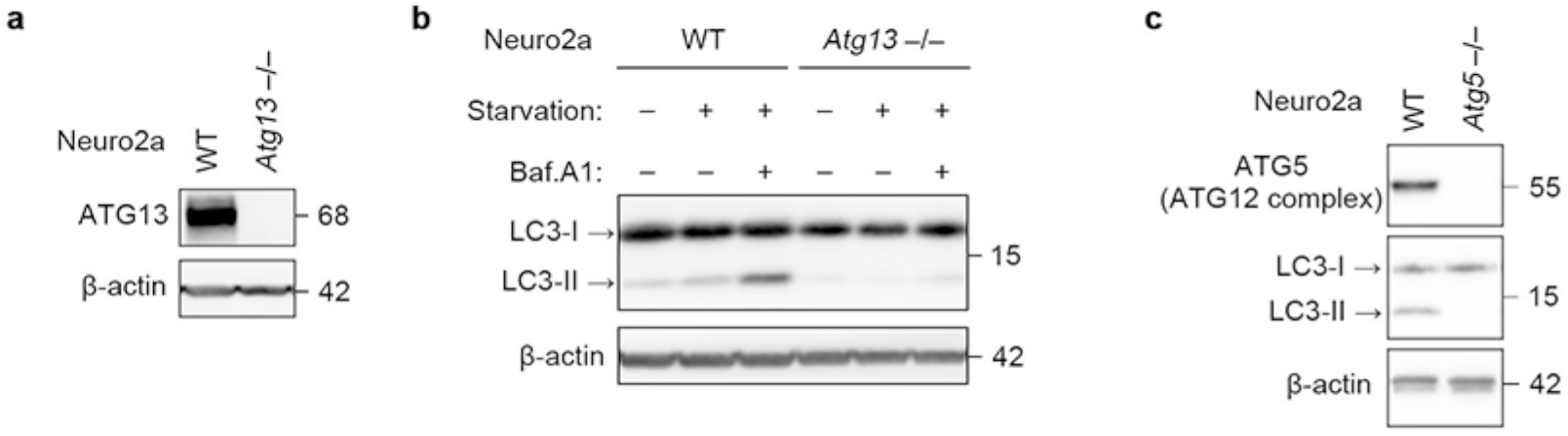
The SIDT2-dependent protein degradation pathway is independent of macroautophagy. (a) Immunoblotting of ATG13 in *Atg13*-KO N2a cells. (b) Immunoblotting of LC3 in WT and *Atg13*-KO N2a cells cultured in normal or starvation medium with or without bafilomycin A1, indicating that macroautophagy is abrogated in the *Atg13*-KO cells. (c) Immunoblotting of ATG5 and LC3 in *Atg5*-KO N2a cells. LC3-II was not detectable in *Atg5*-KO cells, indicating impaired macroautophagy.

**Extended Data Fig. 5.**
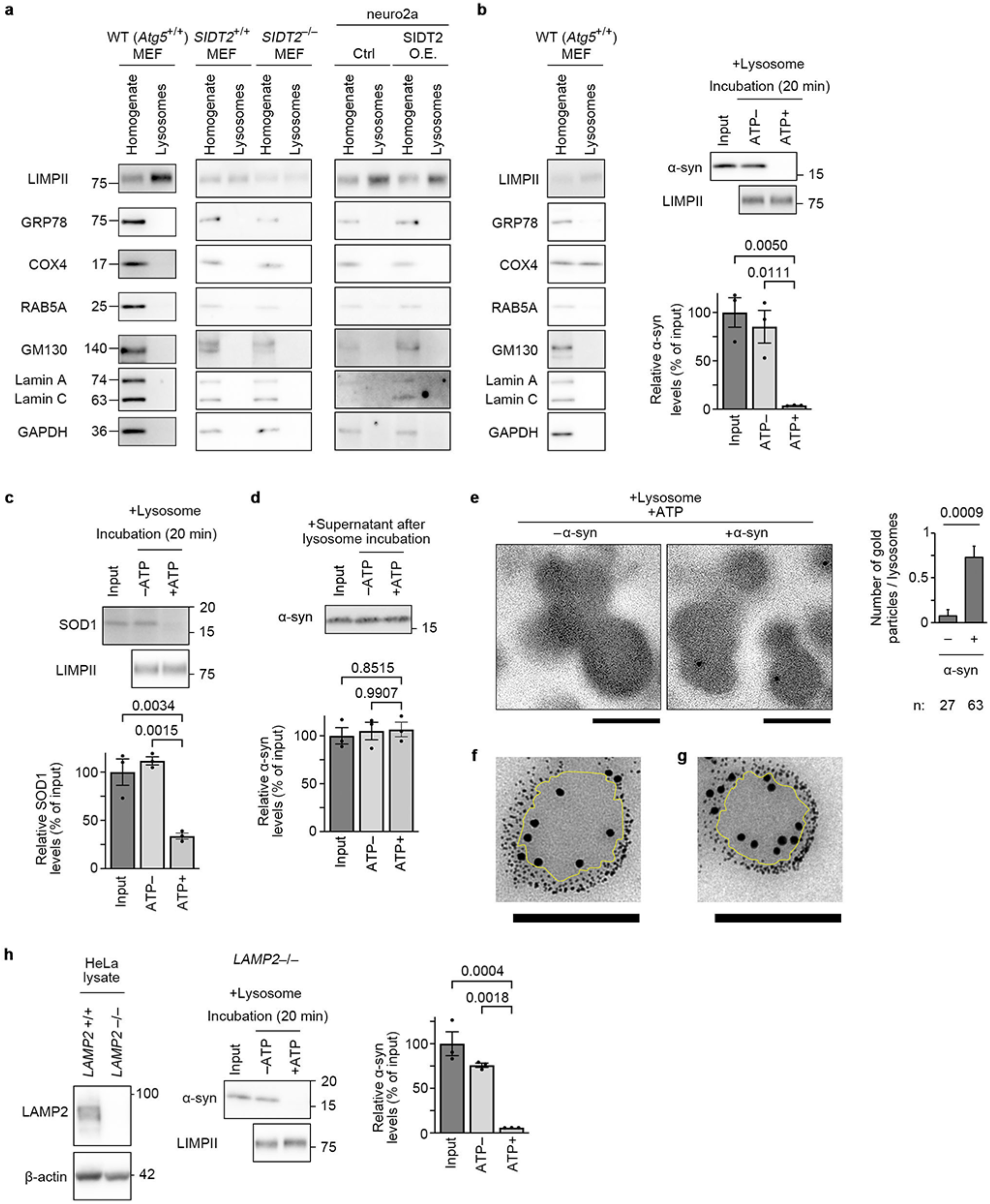
Characterization of lysosomal preparations and lysosome-dependent degradation of cytosolic proteins. (a) Assessment of the purity of lysosomal preparations. Homogenates and lysosomal preparations were prepared from indicated cells and analysed by immunoblotting. LIMPII was used as a lysosomal marker. GRP78, COX4, RAB5A, GM130, lamin A/C, and GAPDH were used as markers for the ER, mitochondria, early endosomes, Golgi, nucleus, and cytosol, respectively. (b) Validation of lysosomal preparations prepared with an alternative method (Lysosome Purification Kit, BioVision). Homogenates and lysosomal preparations from MEFs were analysed using the same organelle markers as in (a). Right panel: α-synuclein was incubated with lysosomes prepared using the BioVision kit in the presence or absence of ATP (n = 3). (c, d) Recombinant cytosolic proteins were incubated with lysosomes to assess ATP-dependent degradation using SOD1 (c) or α-synuclein (d) as a substrate. As a control, α-synuclein was incubated in the supernatant after lysosome pre-incubation with or without ATP (d). No degradation was observed, suggesting that lysosomes remained intact in the lysosomal preparation assays. (e–g) Immunoelectron microscopic observations of α-synuclein protein uptake into lysosomes. Post-embedded immunoelectron microscopy images of α-synuclein protein in lysosomal preparations co-incubated with (e–g) or without (e) recombinant α-synuclein protein in the presence of ATP (e and g, anti-α-synuclein antibody, Abcam; f, anti-α-synuclein antibody, Cell Signaling Technology). Scale bars represent 200 nm. First, 5 nm gold particles were co-incubated with lysosomes and α-synuclein protein in the presence of ATP, and this was followed by immunoelectron microscopy against α-synuclein protein using 15 nm gold particles (f, g). While 5 nm gold particles were adsorbed on the surface of lysosomes, 15 nm gold particles indicating α-synuclein protein were seen in the luminal area of the lysosomes. The yellow line indicates the approximate delimitation between the inside and outside of lysosomes. (h) Left panel: Immunoblotting confirms the loss of LAMP2 expression in *LAMP2*^−/–^ HeLa cells. Right panel: α-synuclein was incubated with lysosomes derived from *LAMP2*^−/–^ HeLa cells under the conditions indicated (n = 3).

**Extended Data Fig. 6.**
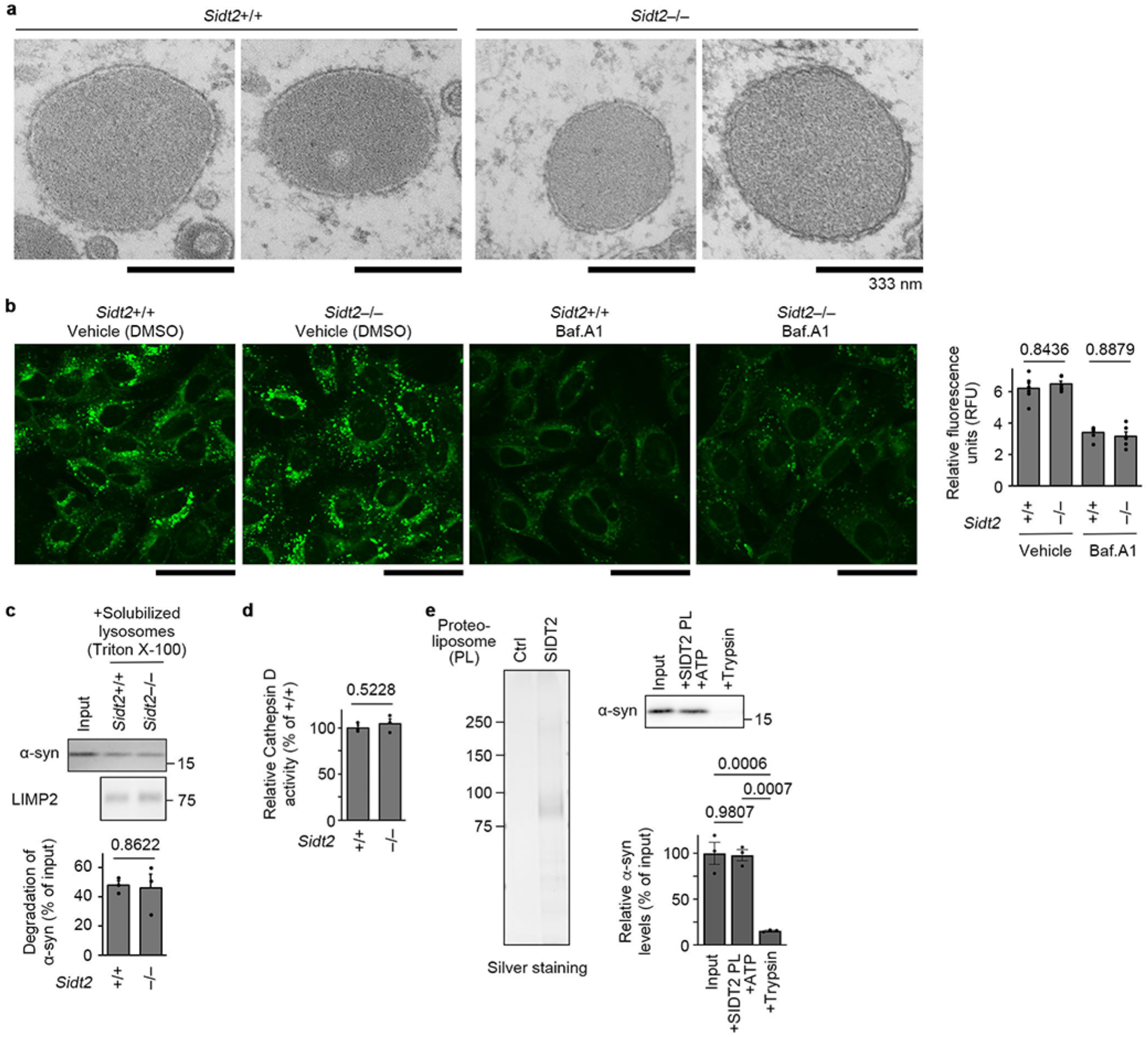
SIDT2 deficiency does not affect lysosomal morphology, pH, or protease activity, and SIDT2 itself shows no detectable proteolytic activity. (a) Electron microscopy images of lysosomes in the indicated cells. No apparent morphological abnormalities were observed in SIDT2-deficient lysosomes. (b) Acidity of lysosomes in *Sidt2^+/+^*and *Sidt2^−/–^* MEFs treated with bafilomycin A1 or DMSO (n = 6). Scale bars = 50 µm. (c) Degradation of α-synuclein protein incubated with dissolved lysosomal preparations derived from *Sidt2^+/+^* or *Sidt2^−/–^* MEFs (n = 3). (d) Cathepsin D activity measured in cell lysates from indicated cells (n = 3). SIDT2 deficiency did not affect cathepsin D activity. The assay was performed using the Cathepsin D Activity Fluorometric Assay Kit (BioVision, #K143-100) according to the manufacturer’s instructions. (e) SIDT2 does not possess intrinsic proteolytic activity. Left: Silver staining of control liposomes and SIDT2 proteoliposomes. Right top: Immunoblotting of α-synuclein after incubation with SIDT2 proteoliposomes in the presence of ATP, or with trypsin. Right bottom: Quantification of α-synuclein levels after incubation (n = 3).

**Extended Data Fig. 7.**
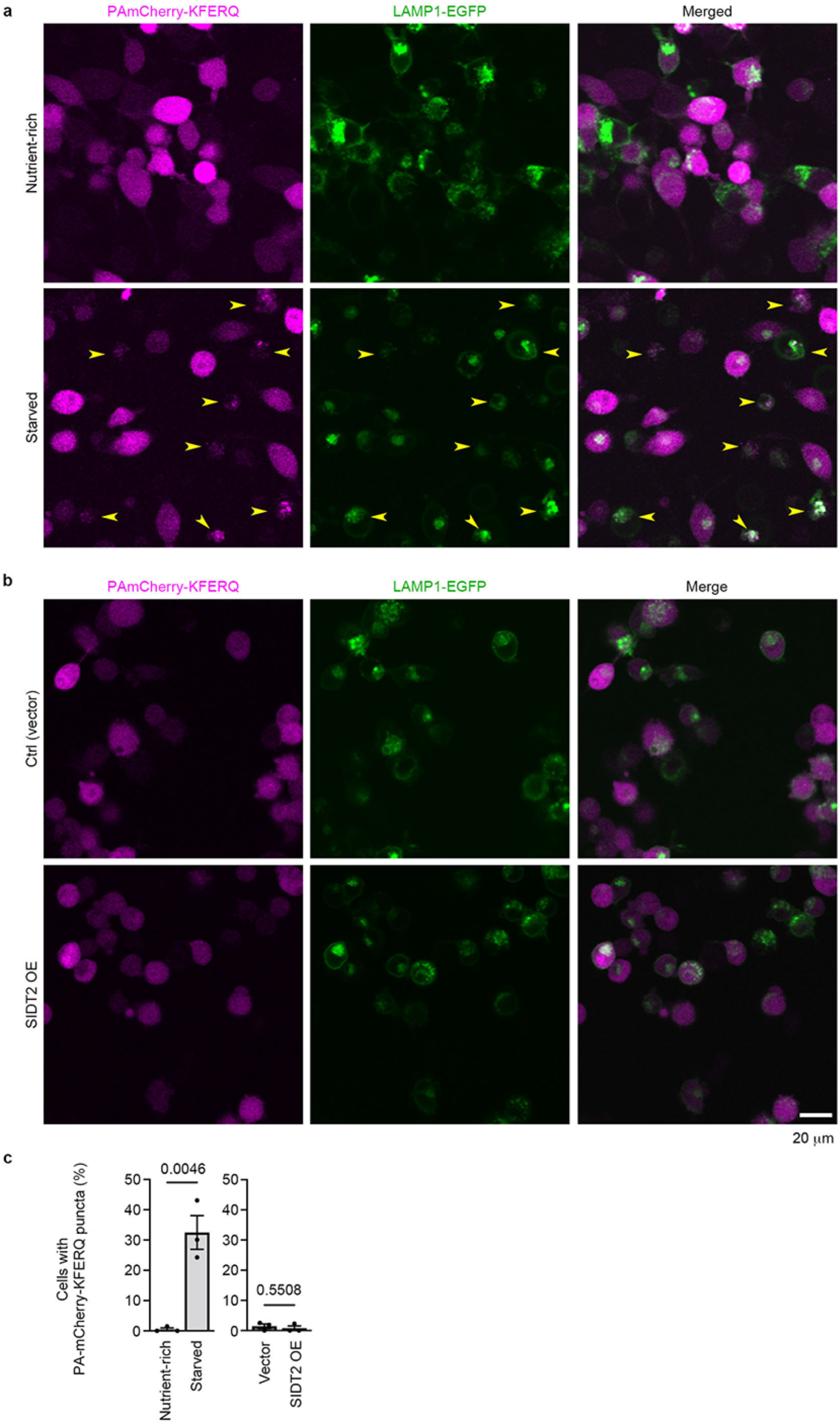
SIDT2 overexpression does not affect formation of chaperone-mediated autophagy (CMA) reporter puncta. (a) Representative fluorescence images showing the localization of PAmCherry–KFERQ, a well-established CMA reporter, and LAMP1–EGFP in N2a cells under nutrient-rich or starvation conditions (24 h). Starvation induced the formation of PAmCherry–KFERQ puncta that colocalize with lysosomes. Yellow arrowheads indicate representative puncta. (b) N2a cells co-expressing PAmCherry–KFERQ, LAMP1–EGFP, and either SIDT2 or empty vector were imaged. (c) Quantification of cells with PAmCherry–KFERQ puncta (n = 3).

**Extended Data Fig. 8.**
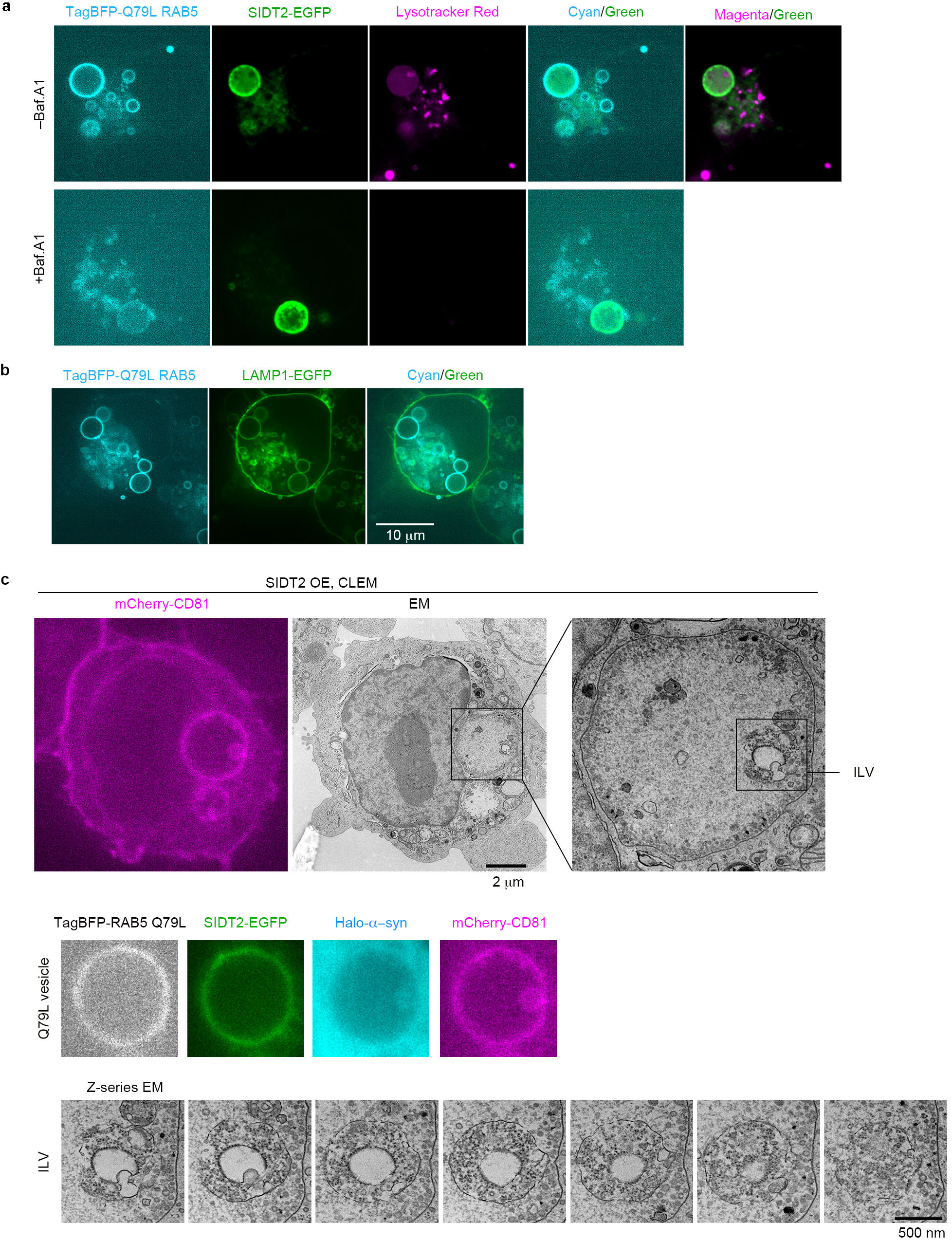
Characterization of Q79L RAB5-induced enlarged endolysosomes and their analysis by CLEM. (a) N2a cells overexpressing TagBFP–Q79L Rab5 and SIDT2–EGFP were treated with or without bafilomycin A1 and stained with LysoTracker. LysoTracker signals were detected in enlarged endolysosomes positive for both Q79L RAB5 and SIDT2, and were abolished by bafilomycin A1 treatment, indicating acidification of these vesicles. (b) Co-overexpression of Q79L RAB5 and LAMP1–EGFP confirmed that the enlarged vesicles were positive for the lysosomal membrane marker LAMP1. (c) CLEM analysis using an enlarged endolysosome positive for both Q79L RAB5 and SIDT2. From a field distinct from that of Fig. 3f.

**Extended Data Fig. 9.**
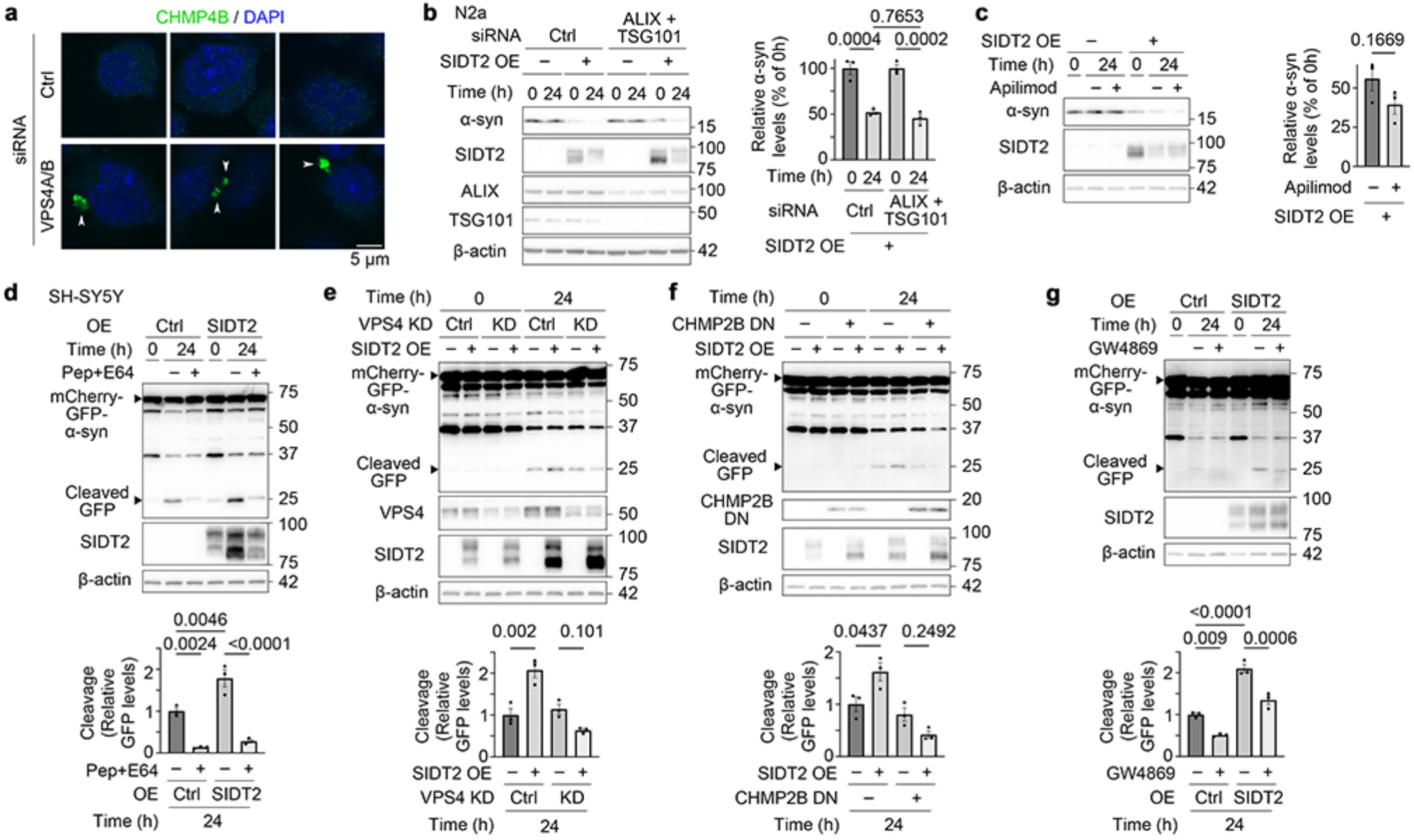
Effects of ESCRT and nSMase perturbation on SIDT2-mediated protein degradation. (a) Accumulation of CHMP4B-positive puncta upon VPS4 knockdown. Representative immunofluorescence images of CHMP4B in N2a cells transfected with control siRNA or siRNAs targeting VPS4A and VPS4B. In control cells, CHMP4B was diffusely distributed, while VPS4 knockdown induced its accumulation in cytoplasmic puncta (arrowheads). Three representative cells are shown for each condition. The accumulation of CHMP4B puncta indicates impaired ESCRT-III disassembly due to loss of VPS4 activity^56^. (b, c) Degradation of α-synuclein protein in N2a cells overexpressing SIDT2, treated with siRNAs against ALIX and TSG101 (b) or the PIKfyve inhibitor apilimod (c). SIDT2-dependent protein degradation was not inhibited. (d–g) SIDT2-dependent protein degradation in SH-SY5Y cells assessed by GFP cleavage assay. Treatment with lysosomal protease inhibitors (E64d and pepstatin A, 50 µg/mL each) suppressed GFP cleavage (d). Knockdown of VPS4A and VPS4B (e), expression of dominant-negative CHMP2B (f), or treatment with GW4869 (g) also impaired SIDT2-dependent enhancement. However, as GW4869 affected basal levels, the dependency on nSMase is likely to be limited.

**Extended Data Fig. 10.**
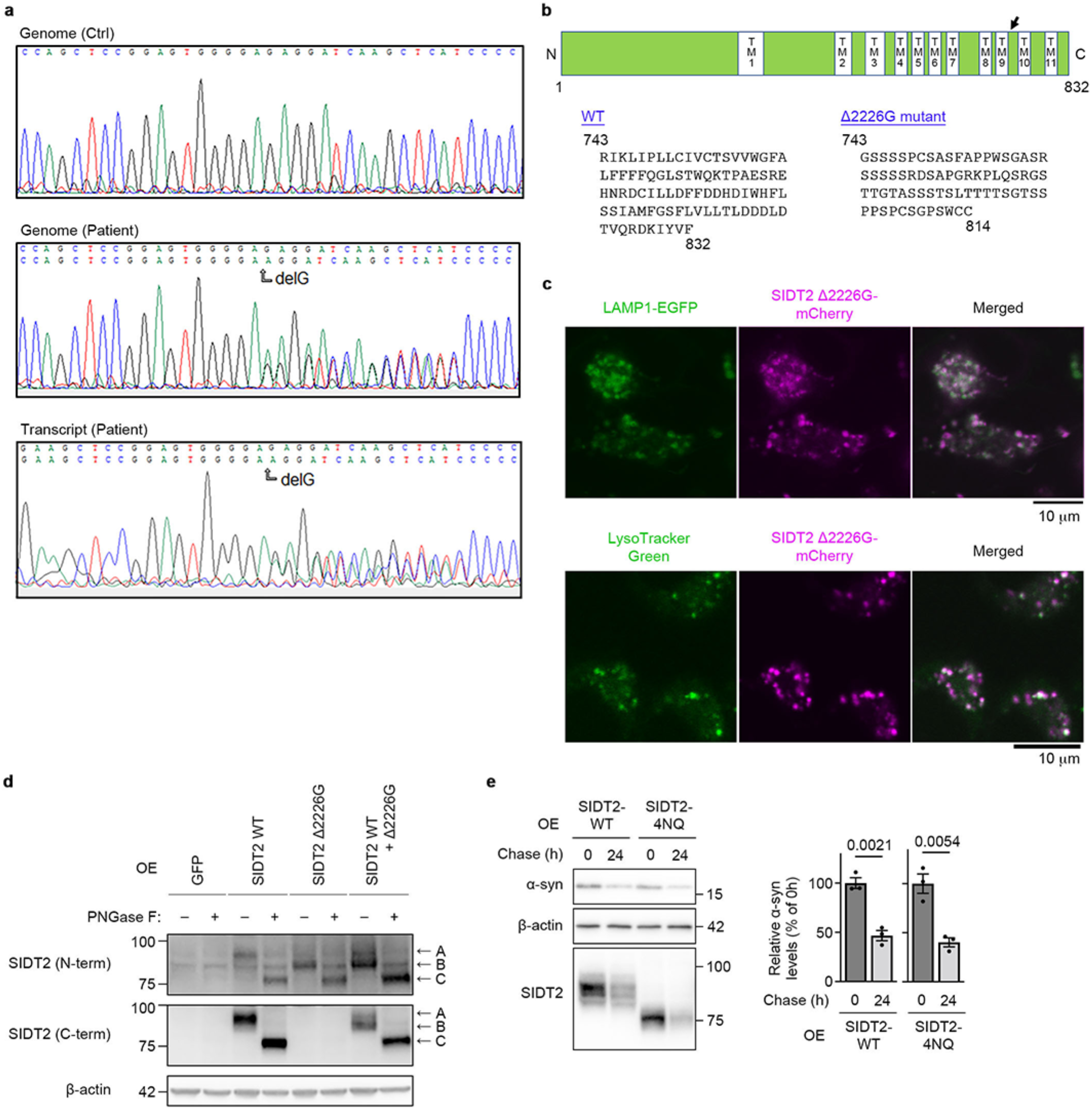
Characterization of rimmed vacuolar myopathy patient harbouring a Δ2226G mutation in SIDT2 and the mutant SIDT2. (a) Sanger sequencing chromatogram of a control biopsy and the proband showing the heterozygous Δ2226G frameshift mutation in *SIDT2,* and direct sequencing chromatogram of the patient’s *SIDT2* transcript. This variant was not listed in any of the following public databases as of May 2026: dbSNP, 1000 Genomes, gnomAD, the Human Gene Mutation Database, and ClinVar. (b) A schematic of SIDT2 protein and sequence of additional aberrant amino acid residues produced by the Δ2226G mutation in SIDT2. The arrow indicates the site of mutation in the patient. (c) Localization of SIDT2(Δ2226G)–mCherry in N2a cells co-transfected with LAMP1–EGFP (upper panels) or stained with LysoTracker Green (lower panels). The mutant protein is correctly targeted to lysosomes. (d) Glycosylation of SIDT2–WT and SIDT2–Δ2226G. (e) Degradation of α-synuclein protein in N2a cells overexpressing SIDT2–WT or SIDT2–4NQ, in which all predicted N-glycosylation sites were mutated. The 4NQ mutant promoted protein degradation to a similar extent as WT, indicating that glycosylation is not required for SIDT2-mediated protein degradation.

**Extended Data Fig. 11.**
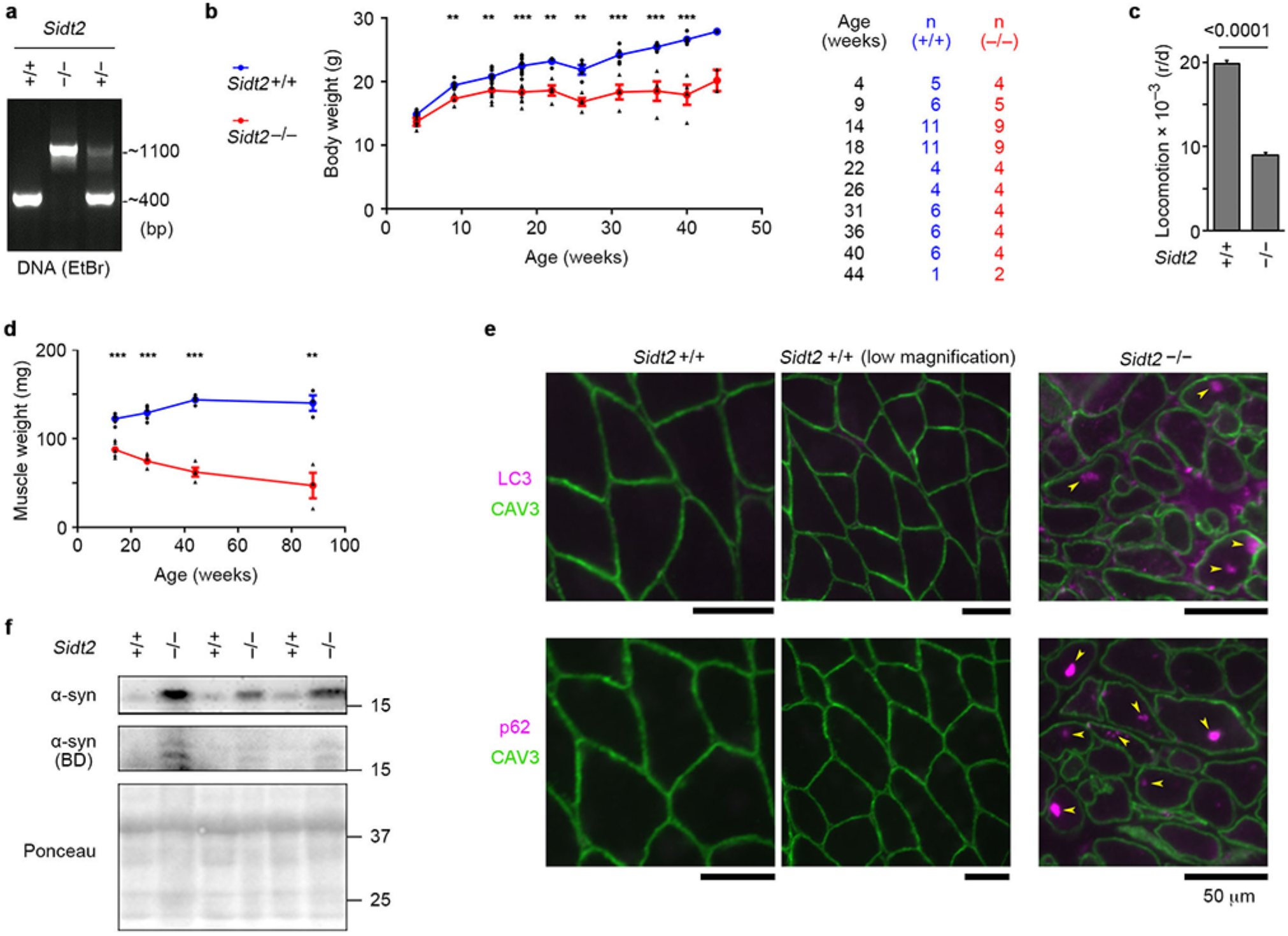
Characterization of *Sidt2* knockout mice. (a) Genotyping of *Sidt2* by PCR showing the WT allele-specific band at approximately 400 bp, and the KO allele-specific band at approximately 1,100 bp. (b) Growth curves of *Sidt2*^+/+^ and *Sidt2^−^*^/*–*^ mice. (c) Locomotor activities of *Sidt2*^+/+^ and *Sidt2^−^*^/*–*^ mice (+/+, n = 154 from 11 mice; ^−^/^−^, n = 126 from 9 mice). (d) Changes in weight of the gastrocnemius of *Sidt2*^+/+^ and *Sidt2^−^*^/*–*^ mice (n = 3–5). (e) Immunofluorescence analysis of skeletal muscle sections from *Sidt2*^+/+^ and *Sidt2*^−/–^ mice. Yellow arrowheads indicate LC3- or p62-positive deposits. The images provide additional examples supporting the observations in Fig. 5g. (f) Immunoblot analysis of α-synuclein in skeletal muscles of *Sidt2*^+/+^ and *Sidt2*^−/–^ mice using an anti-α-synuclein antibody (BD Transduction Laboratories, 610787). Another α-synuclein blot and the Ponceau staining are the same as in Fig. 5h. In (b) and (d) Results are the mean ± s.d. \*\**P* < 0.001, \*\*\**P* < 0.0001.

**Extended Data Fig. 12.**
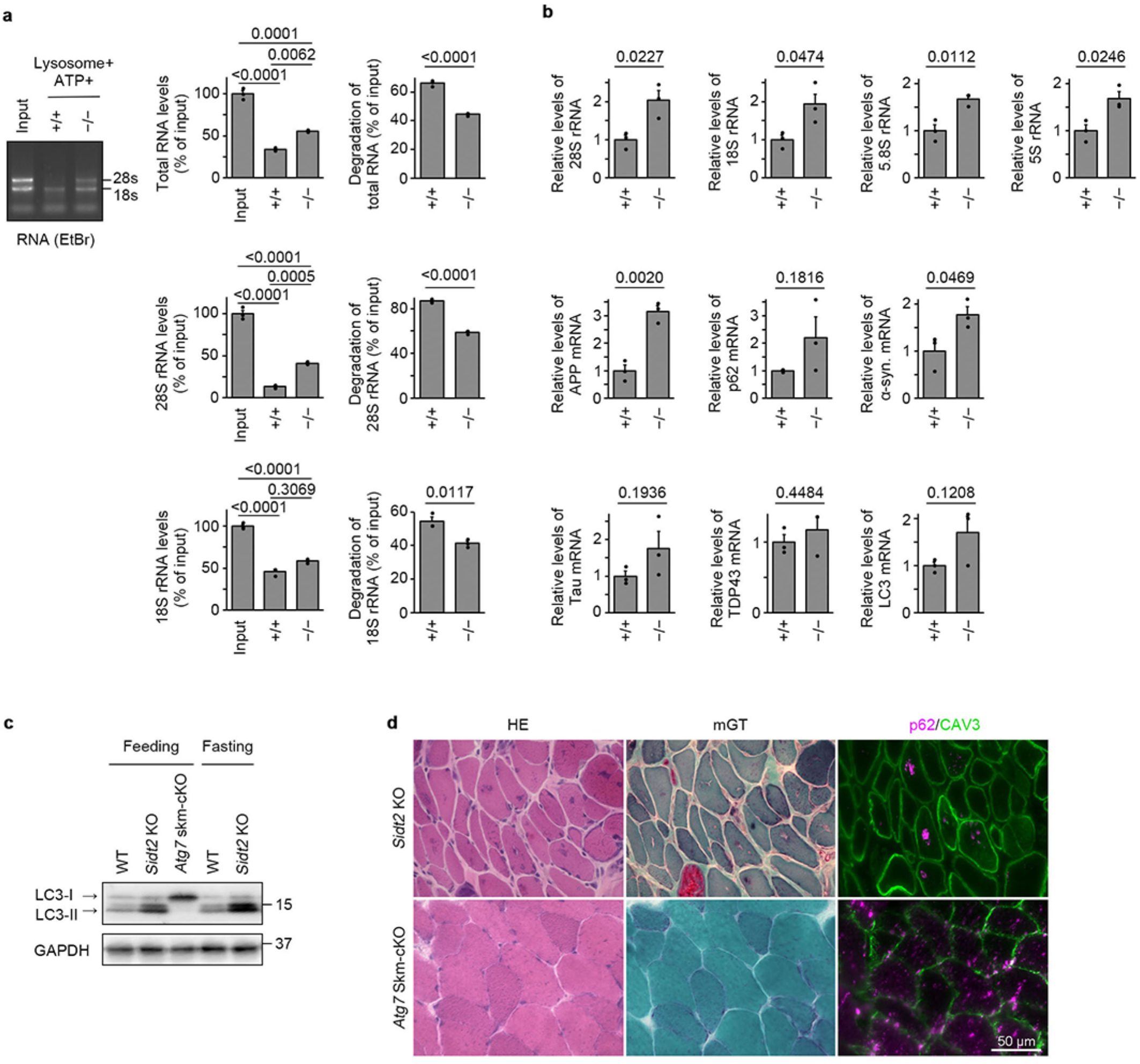
Impaired RNA degradation in SIDT2-deficient muscle and lack of rimmed vacuole formation in macroautophagy-deficient muscle. (a) Degradation of RNA by lysosomal preparations derived from *Sidt2*^+/+^ or *Sidt2^−^*^/*–*^ mouse skeletal muscles (n = 3). (b) Levels of RNA in gastrocnemius of *Sidt2*^+/+^ or *Sidt2^−^*^/*–*^ mice (n = 3). (c) Immunoblotting of LC3 in feeding and fasting *Sidt2*^+/+^ and *Sidt2^−^*^/*–*^ mouse skeletal muscles. (d) Comparison of skeletal muscle pathology of *Sidt2^−^*^/*–*^ mice and *Atg7* skeletal muscle conditional KO mice. Although atrophy was also observed, rimmed vacuoles were not observed, and p62 deposits were more diffusely observed in the macroautophagy-deficient muscles, compared with those observed in *Sidt2^−^*^/*–*^muscles.

**Extended Data Fig. 13.**
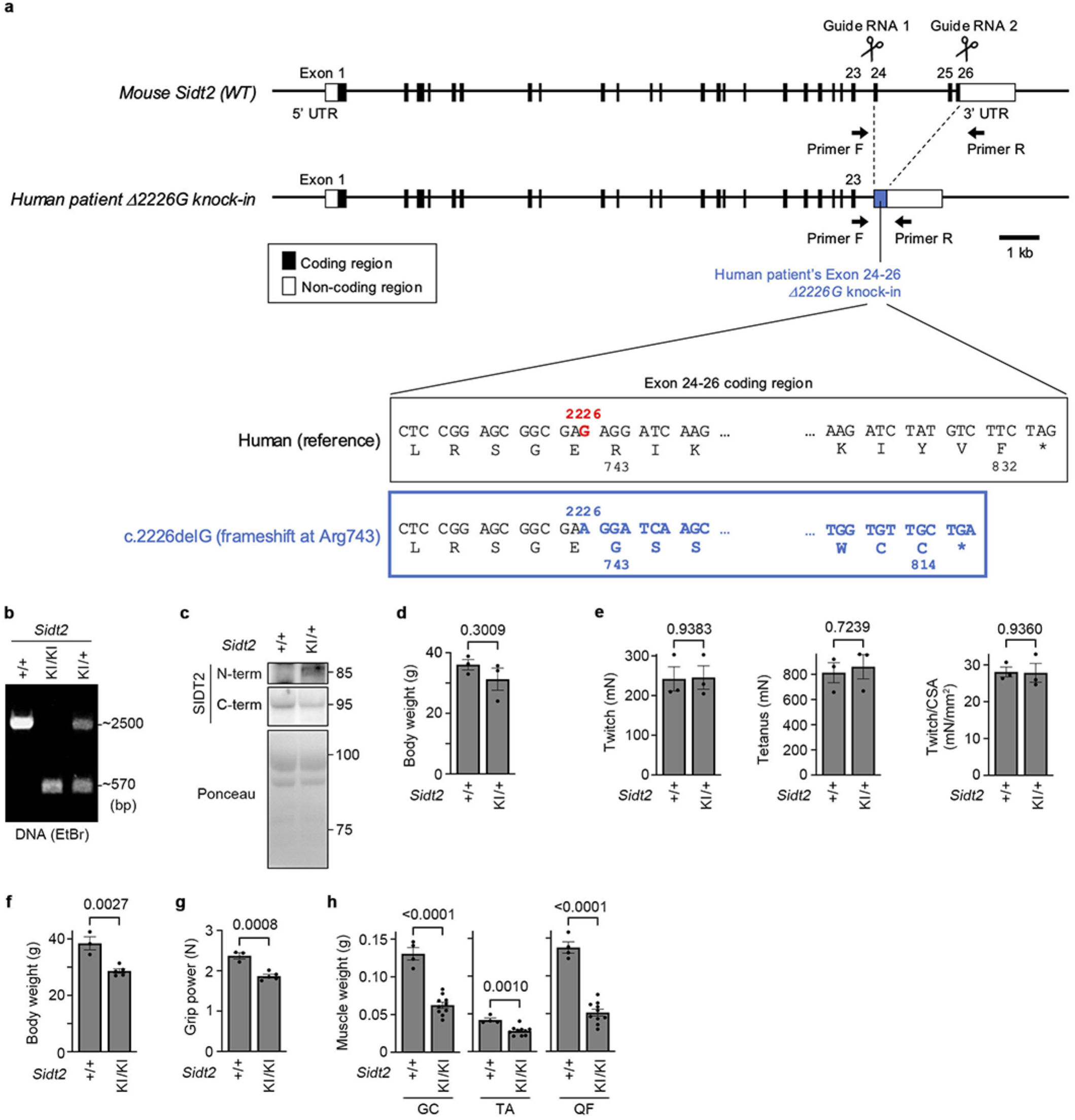
Characterization of *Sidt2* knock-in mice expressing SIDT2 with an equivalent amino acid change to the Δ2226G patient mutation. (a) Generation of knock-in (KI) mice carrying the human c.2226delG (Arg743 frameshift) mutation by CRISPR/Cas9. Targeting strategy to delete the mouse *Sidt2* exon 24–26 and insert the human patient-derived frameshifted exon 24–26 sequence is schematically outlined. Guide RNA 1 and Guide RNA 2 were selected based on their proximity to the deletion sites and minimal off-target effects. The 2226G, deleted in a human patient, is marked with red colour, and the resulting frameshift at Arg743 is shown in blue colour. 90 amino acid residues are replaced with 72 amino acid residues of an aberrant sequence. For genotyping, Primer F and Primer R, indicated by black arrows, are used. (b) Genotyping of *Sidt2* KI mice by PCR. WT allele-specific band: ∼2500 bp; KI allele-specific band: ∼570 bp. (c) Immunoblotting of skeletal muscle lysates from *Sidt2*^+/+^ and *Sidt2*^KI/+^ mice. Ponceau S staining is shown as a loading control. (d) Body weight of *Sidt2*^+/+^ and *Sidt2*^KI/+^ mice at 104 weeks of age (n = 3). (e) Muscle contraction analysis of gastrocnemius muscle from *Sidt2*^+/+^ and *Sidt2*^KI/+^ mice at 104 weeks of age. Twitch force (left), tetanus force (middle), and specific twitch force normalized by cross-sectional area (CSA) (right) (n = 3). (f, g) Body weight and (f) forelimb grip power (g) of *Sidt2*^+/+^ and *Sidt2*^KI/KI^ mice at 94 weeks of age (+/+, n = 3; KI/KI, n = 5). (h) Muscle weight of gastrocnemius (GC), tibialis anterior (TA), and quadriceps femoris (QF) in *Sidt2*^+/+^ and *Sidt2*^KI/KI^ mice at 112–117 weeks of age (+/+, n = 4; KI/KI, n = 10).

**Extended Data Fig. 14.**
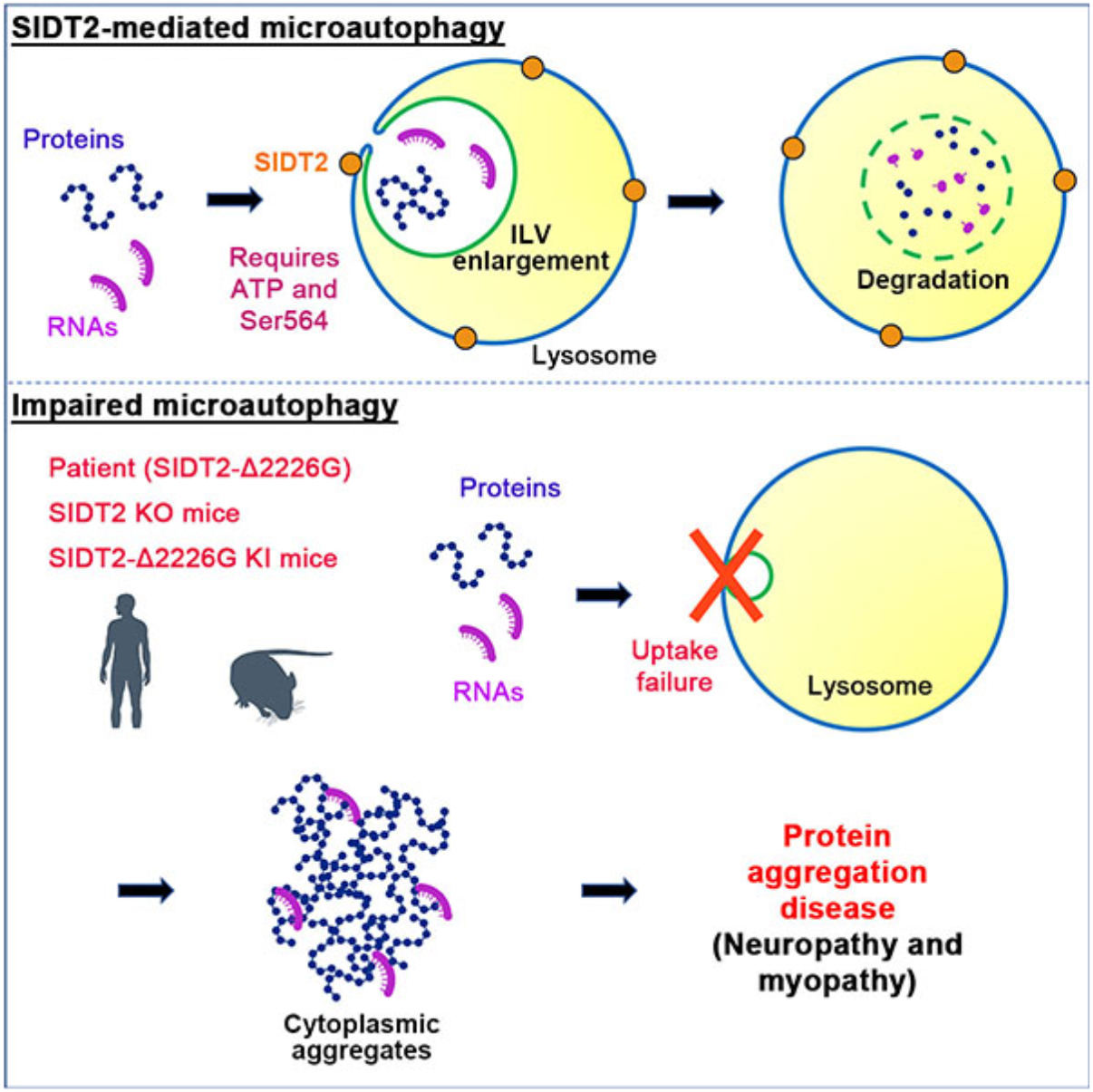
A schematic of SIDT2-mediated microautophagy dysfunction leading to protein aggregation disease. SIDT2-mediated microautophagy maintains cellular homeostasis by degrading cytosolic protein and RNA (upper panel). Impairment of this pathway caused by mutation or deficiency in SIDT2 results in an accumulation of cytoplasmic aggregates and deposits, eventually causing neuromuscular dysfunction and protein aggregation disease (neuropathy and myopathy with rimmed vacuoles) (lower panel).

## Legends for supplementary movies

**Movie S1**. Z-series EM images of an ILV in an enlarged endolysosome of SIDT2-overexpressing cells (related to Fig. 3g).

**Movie S2**. Z-series EM images of an ILV in an enlarged endolysosome of SIDT2-overexpressing cells (related to Extended Data Fig. 8c).

